# The allelopathic vitamin B1 antagonist bacimethrin impacts microbial gene expression in a hypereutrophic watershed dominated by cyanobacterial blooms

**DOI:** 10.1101/2025.07.29.667554

**Authors:** Kelly C. Shannon, Frederick S. Colwell, Byron C. Crump, Elizabeth Brennan, Gillian St. John, Robin Gould, Christopher Hartzell, McKenzie Wasley, Christie Nichols, Clifford E. Kraft, Christopher P. Suffridge

## Abstract

Freshwater cyanobacterial harmful algal blooms (cyanoHABs), often dominated by *Aphanizomenon*, *Dolichospermum*, and *Microcystis,* are intensifying in eutrophic watersheds globally. A potential control on bacterioplankton dynamics in these systems is the availability of the essential metabolic cofactor thiamin (vitamin B_1_) and presence of the allelopathic thiamin antagonist, bacimethrin, that competitively inhibits thiamin-requiring enzymes. We examined dissolved concentrations of thiamin chemical congeners and bacimethrin, 16S-amplicon based microbiome compositions, prokaryotic mRNA-based metatranscriptomes, and reference genomes in hypereutrophic Upper Klamath Basin before and during seasonal cyanoHABs. Our objective was to connect bacterioplankton community compositions and gene expression patterns with thiamin congener and bacimethrin availability under different cyanoHAB conditions.

Bacimethrin was present in all samples at nearly equimolar concentrations to the thiamin precursor, HMP, suggesting that similar mechanisms influence the availability of both compounds. Additionally, bacimethrin concentrations were positively correlated with cyanoHAB species abundance (cells mL^-1^) and the expression of microbial thiamin biosynthesis genes.

Samples with high cyanoHAB abundances displayed elevated transcription of genes for thiamin biosynthesis, the pentose phosphate pathway, and photosynthesis. Bacterioplankton unable to synthesize thiamin and thus vulnerable to bacimethrin allelopathy, such as *Limnohabitans* spp., showed reduced gene expression when cyanoHAB abundances were high. Reference genomes of cyanoHAB and many picocyanobacteria strains contained complete thiamin biosynthesis gene pathways, implicating these taxa as major thiamin sources. These results suggest that bacimethrin provides a competitive advantage to bacterioplankton that do not require exogenous vitamin B1 by eliminating the risk of bacimethrin uptake with vitamin B1 transporters, potentially facilitating cyanoHAB dominance in Upper Klamath Basin and broader eutrophic watersheds.

## Introduction

Discovering mechanistic drivers of microbial interactions is a central goal in microbial ecology, particularly in aquatic ecosystems where diverse bacterioplankton communities mediate regional to global scale biogeochemistry, shaping ecosystem function at large scales [1]. Recent advances in meta-omics have greatly accelerated this effort by integrating environmental data, including dissolved metabolite profiles, into large scale correlative analyses of microbial community composition and function [2–4]. This approach enables more direct inference of ecological processes and facilitates the generation hypotheses about the underlying mechanisms driving correlative trends. In this study, we sought to describe bacterioplankton interactions and biogeochemistry in hypereutrophic freshwater environments by combining dissolved measurements of the essential metabolite thiamin, its chemically related congeners, and its toxic antagonists, with microbial community data.

The complex biosynthesis and exchange of thiamin, its congeners, and bacimethrin (thiamin and related compounds; TRCs hereafter) provide an ideal framework for examining microbial interactions. This essential coenzyme is required across all domains of life and microbial communities produce the bulk of thiamin for aquatic food webs [5, 6]. However, most microbes lack the complete thiamin biosynthesis pathway and must salvage TRCs from the dissolved pool [7–9]. As such, it has been shown that dissolved TRC availability can structure bacterioplankton communities [10, 11]. Thiamin is a coenzyme that, in its active form of thiamin pyrophosphate (TPP), is required by enzymes key to both catabolic and anabolic carbon metabolism including the tricarboxylic acid cycle, the Calvin cycle, branched chain amino acid biosynthesis, and the pentose phosphate pathway [9].

Microorganisms range in their ability to synthesize thiamin *de novo*, and many bacterioplankton species are thiamin auxotrophs, lacking the full genomic pathway for producing thiamin [7]. Auxotrophic bacterioplankton must import exogenous thiamin or thiamin precursors, including the pyrimidine 4-amino-5 hydroxymethyl-2-methylpyrimidine (HMP) and the thiazole 5-(2-hydroxyethyl)-4-methyl-1,3-thiazole-2-carboxylic acid (cHET) [12], to meet cellular thiamin requirements [7], with pyrimidine (HMP) auxotrophies being the most prevalent [7, 13–15]. Thiamin is also abiotically degraded into 4-amino-5-aminomethyl-2-methylpyrimidine (AmMP; pyrimidine degradation product) and 4-methyl-5-thiazoleethanol (HET; thiazole degradation product), which some organisms can recycle to synthesize thiamin [8, 13, 16, 17]. In aquatic ecosystems, the net exchange of TRCs between producers (typically prototrophs that have the full *de novo* thiamin biosynthesis pathway) and consumers (typically auxotrophs) controls dissolved TRC concentrations [13, 18] and – when coupled with microbial community data – provides a framework for understanding thiamin acquisition by microorganisms [19].

Bacimethrin, an understudied thiamin analog with allelopathic properties, has been identified as a microbial secondary metabolite produced by cultured soil bacteria [20–22] and, likely, microbes in other environments. As a toxic analog of HMP [20, 21], bacimethrin targets thiamin auxotrophs that lack HMP biosynthesis genes by interrupting proper thiamin biosynthesis. The incorporation of bacimethrin rather than HMP into the pyrimidine branch of the thiamin biosynthesis pathway leads to the formation of methoxy-thiamin pyrophosphate [23, 24], a dysfunctional coenzyme that competitively inhibits thiamin-dependent enzymes, thereby disrupting essential metabolic functions. Although extensively characterized in laboratory cultures, the ecological role of bacimethrin in natural environments remains poorly understood. In aquatic ecosystems, bacimethrin may alter bacterioplankton community structure by mimicking HMP and interfering with its uptake via HMP transporters [25] and use via thiamin biosynthesis enzymes [26].

Recent evidence shows that *Microcystis* spp., which form extensive cyanobacterial harmful algal blooms (cyanoHABs), can synthesize bacimethrin [27]. As thiamin prototrophs [9, 28, 29], *Microcystis* spp. and related cyanobacteria are likely resistant to bacimethrin allelopathy, and cyanoHAB-produced bacimethrin may confer a selective advantage to prototrophs by competitively inhibiting thiamin or HMP-utilizing enzymes in auxotrophic taxa. This mechanism could facilitate cyanoHAB niche expansion and persistence through disruption of native HMP auxotrophic microbial assemblages.

This potential niche expansion by cyanoHABs – via the competitive exclusion of bacimethrin-sensitive (must import exogenous HMP) bacterioplankton – is of substantial global importance, as the frequency and severity of cyanobacterial harmful algal blooms in freshwater ecosystems has increased [30] alongside anthropogenic stressors such as increased water temperature, nutrient pollution, agricultural and industrial activity, and urbanization [31, 32].

CyanoHABs magnify eutrophication by producing toxins (cyanotoxins) and altering aquatic food web stability [33, 34], with impacts [35] that cost billions of dollars annually to manage due to their detrimental impacts on freshwater ecosystems [30, 36]. Upper Klamath Basin, OR (UKB) provides an ideal environmental setting to investigate the impacts of bacimethrin-based allelopathy on microbial ecology. CyanoHABs have been extensively studied in the UKB [37, 38], and sediment cores from Upper Klamath Lake (the lake only rather than the whole basin; UKL) indicate that increases in cyanoHAB species abundance in the early 20^th^ century correspond with an intensification of eutrophication driven by increased N and phosphorus (P) inputs from previously drained marshlands and agricultural activity [39]. Over decadal times scales, cyanoHAB species have gradually replaced diatoms in UKL during summer months [40]. Contemporary UKL cyanoHABs, which can also impact other UKB habitats (rivers and reservoirs), are typically initiated by blooms of filamentous and diazotrophic (can fix atmospheric nitrogen; N_2_) *Aphanizomenon* and *Dolichospermum* spp. in spring [41]. These cyanoHAB species grow rapidly during the early summer and reach peak biomass in June or July, corresponding to an annual peak and subsequent rapid decline in water pH, dissolved oxygen (DO), and primary production [41]. Blooms of *Microcystis* generally follow filamentous cyanoHAB species and have similarly been linked to habitat degradation in UKL [42, 43].

We investigated impacts of the availability of TRCs (thiamin, HET, cHET, AmMP, HMP, and bacimethrin; all dissolved) on microbial communities and cyanoHABs in the UKB by pairing measurements of TRC concentrations with 16S rRNA gene-based analysis of microbiomes, metatranscriptomes, and reference genome assemblies. Samples were collected prior to (May) and during (August) a seasonal cyanobacteria bloom in UKL and its springs, tributaries, outflow, and surrounding reservoirs. We sought to (1) investigate the environmental distributions of TRCs across diverse UKB habitats with varying cyanoHAB conditions, (2) identify UKB habitats, cyanoHAB-related seasons (prior to and during the time when cyanoHABs form; spring and summertime), and conditions associated with the greatest potential for bacimethrin allelopathy to bacterioplankton, and (3) link patterns of bacterioplankton gene expression and thiamin biosynthesis gene activity with TRC concentrations in water samples with contrasting impacts (high & low) of cyanoHABs.

## Materials and Methods

### Sampling procedure, sample site description, and water quality measurements

Microbial and TRC surface water samples were collected May 1-3 and August 28-30, 2023, during daylight hours following previously described methods [18]. Sampling sites were selected to examine variable impacts of cyanoHABs and to contrast lotic (UKL tributaries and outflow) and lentic (UKL and reservoirs) systems (Figure 1A; see Supp. Methods for sample site description). In UKL, *Aphanizomenon* and *Dolichospermum* spp. blooms typically begin in late May and *Microcystis* spp. blooms begin in late July, after which both blooms can persist into late Fall [44]. Thus, samples were binned into pre-bloom (early-May) and bloom (late-August) periods. Publicly available USGS data (https://waterdata.usgs.gov/nwis) for UKL water elevation, Sprague and Williamson River discharge, and Link River pH and DO were accessed and plotted with the dataRetrieval USGS R package [45] (see Supp. Methods).

**Figure 1.**
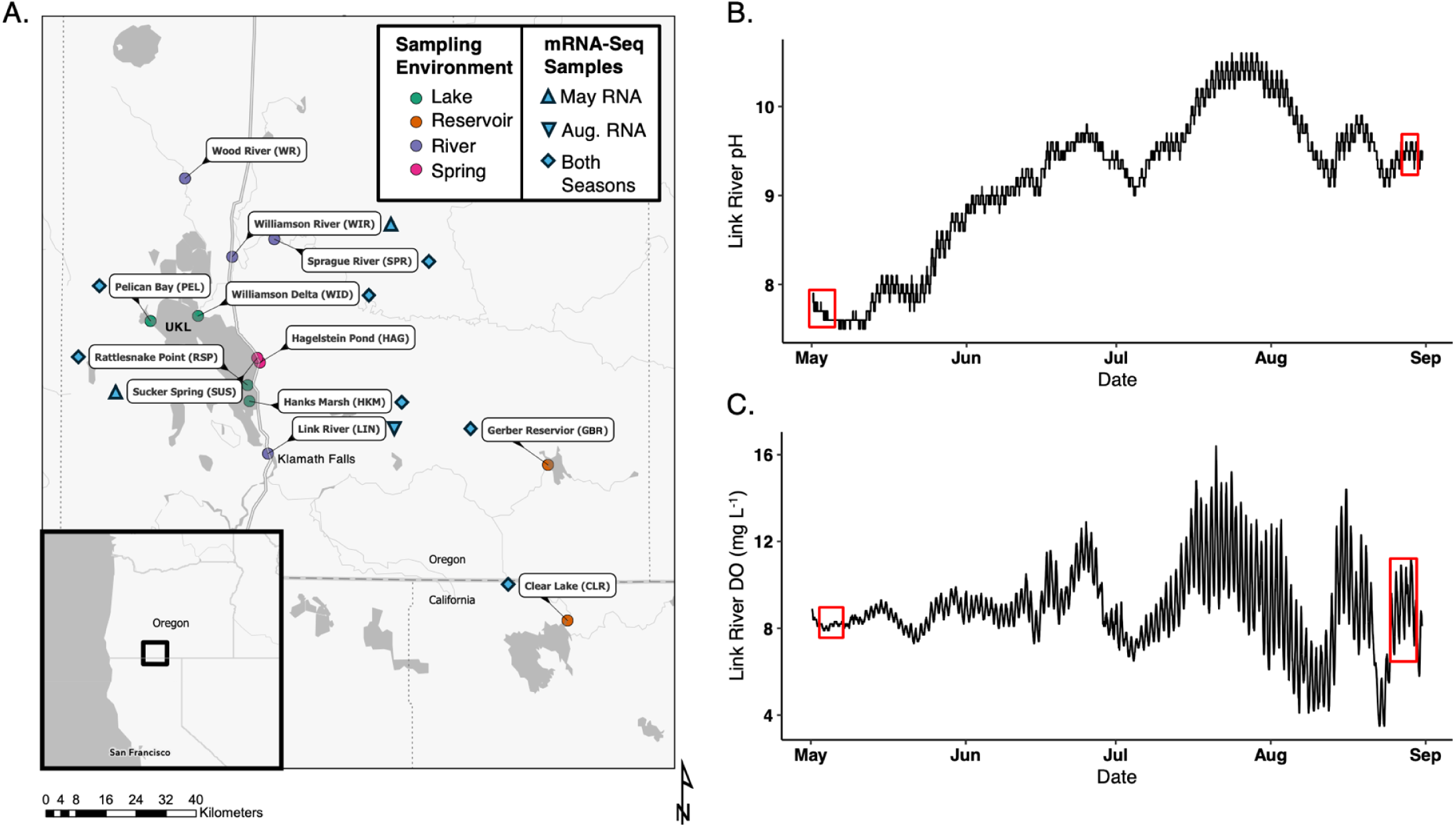
A map of sampling locations and ecological conditions in Upper Klamath Lake between late spring and summer. (A) Sample map showing the locations of sample collection. The environment type is indicated by color of sampling points. Light blue shapes indicate if sufficient RNA was extracted from samples to produce metatranscriptomes and, if so, which seasons are represented by metatranscriptomes at each sample site; May = pre-bloom and Aug. = bloom time periods. Sample abbreviations shown here apply to subsequent figures. Analyses for 16S, dissolved thiamin congener and dissolved bacimethrin were conducted at all locations and time points. USGS continuous water quality measurements of (B) pH, (C) dissolved oxygen measurements at a Link River USGS water quality station (Supp. Methods) were compiled and plotted between May 5^th^ and Sept. 1^st^, 2025. Red boxes indicate the range of sampling dates for DNA, RNA, and TRC measurements.

Samples were collected at ∼0.5 m depth using 1 L amber HDPE bottles (Nalgene). All samples were prefiltered to remove metazoans (100 µm mesh filter) and stored on ice until further processing. Within the same day of sampling, peristaltic filtration at a rate of 30 mL min^−1^ was used to collect cells and particles onto 0.22-µm Sterivex filters (PES membrane, Millipore, Burlington, MA, U.S.A.) and volumes of filtered water were recorded. Immediately following filtration, 1 mL of RNA*later* (Thermo Fisher Scientific) was pipetted directly into Sterivex cartridges, which were then sealed and incubated in the dark for 5 minutes. Following incubation, sealed Sterivex cartridges were flash frozen in liquid nitrogen and immediately placed in a-80 °C freezer until analysis.

Cell-free filtrate was used for TRC and nutrient concentration measurements. TRC samples were collected in acid-washed and methanol-rinsed amber HDPE bottles, acidified with 1 mL of 1 M hydrochloric acid, and stored at-20 °C until analysis at Oregon State University (OSU). Nutrient samples were collected in acid washed glass vials, frozen at-20°C and shipped to the University of California, Santa Barabara Marine Science Institute Analytical Lab for analysis using their flow injection system for nitrate + nitrite, ammonium, and phosphate.

Temperature (°C) and DO (mg L^−1^) were measured with a Thermo Fisher Scientific (Waltham, MA, U.S.A.) Orion multiparameter meter with RDO and temperature sensors. Water quality conditions were predicted using the Kann and Walker (46) surface water elevation model. This model predicts the probability of poor water quality conditions based on surface elevation of the lake, where deviations from normal surface elevations – based on historical data – will predict high likelihoods of poor water quality conditions.

### Dissolved TRC Analysis

Dissolved TRCs were extracted from water samples and analyzed using LC-MS as previously described [18, 27]. Briefly, samples were thawed, pH adjusted to 6.5, and TRCs were extracted using C_18_ resin (Agilent Bondesil HF). TRCs were eluted from the C_18_ resin using 12 mL of methanol, then further concentrated by nitrogen drying to a volume of 250 μL. A 1:1 chloroform liquid phase extraction was used to remove hydrophobic compounds from the sample matrix.

Analysis was conducted using an Applied Biosystems 4000 Q-Trap triple quadrupole mass spectrometer with an ESI interface coupled to a Shimadzu LC-20AD liquid chromatograph. Chromatography and mass spectrometer conditions for all TRCs are described elsewhere [18, 19, 27]. Samples were analyzed in triplicate and were randomized prior to analysis. An internal standard (^13^C-labeled thiamin) was used for quantification. The inclusion of the internal standard allowed for concentrations to be corrected for matrix effects. Analysis was conducted at the OSU Mass Spectrometry Center. A subset of the bacimethrin concentration data (Hanks Marsh and Williamson River sites) (Figure 1A) have been previously published by Mohammad Yazdani, Christopher P. Suffridge (27).

### Microbial DNA and RNA extractions and sequencing

All DNA and RNA extraction and purifications were performed with ZymoBIOMICS (Irvine, California, U.S.A.) DNA/RNA Miniprep Kits in a biosafety hood that was sequentially sterilized with UV radiation, Obliterase (Innovative Scientific Solutions; Dayton, Ohio, U.S.A.), and 70% ethanol (see Supplemental Methods). Manufacturer instructions were followed except for minor changes to the initial bead-beating procedure (see Supp. Methods). RNA samples were further purified and concentrated with ZymoBIOMICS clean and concentrator kits following manufacturer instructions except that the in-column DNase I treatment was performed only once, following the first cleaning step, and that RNA was eluted with 30 µl DNase/RNase-free water. Internal standards were added prior to extraction for DNA absolute abundance measurements (Supp. Methods for full details) [47].

For DNA, PCR was performed with Platinum II Hot-Start polymerase (2x) (Invitrogen; Waltham, Massachusetts, U.S.A.) to amplify the prokaryotic V4 region of the 16S rRNA gene with 2.5 µl of 2 µM 515F-806R primers [48]. 16S amplicons were sequenced with Illumina (San Diego, California, U.S.A.) MiSeq 2×250 bp paired-end high throughput sequencing (HTS) (Supp. Methods for PCR protocol and MiSeq preparation). Purified and concentrated RNA was sent to OSU Center for Quantitative Life Science for Illumina NextSeq 2×100 bp paired-end HTS, following sequencing preparation with the Illumina Ribo-Zero Plus rRNA Depletion Kit. Following this step, 16 of the 24 samples had enough RNA for sequencing (Figure 1A).

### DNA sequencing data bioinformatics and statistics

Statistical analyses were performed in RStudio (v4.2.1) and code for the generation of all DNA and RNA figures can be found on GitHub (Data Availability Statement). 16S rRNA gene sequencing data was demultiplexed and processed following previous methods [18]. Briefly, adaptor trimming and initial quality filtering (sequences with Phred scores <20 were dropped) was performed with TrimGalore (v0.6.6) [49] and read processing to amplicon sequence variants (ASVs) was performed with DADA2 (v1.24.0) [50] following default parameters aside from the inclusion of pseudo pooling during ASV assignment.

To improve comparability between 16S and metatranscriptome taxonomic results, the Genome Taxonomy Database (GTDB; v202)[51] was used for 16S taxonomic annotations other than those for *T. thermophilus*-based absolute abundance calculations (Supp. Methods for detailed *T. thermophilus* steps). All 16S community composition and beta-diversity plots were made with the phyloseq (v1.42.0) and microViz (v0.11.0) packages [52, 53]. Aitchison distance (Euclidean distance between samples after a center-log ratio (CLR) transformation) was used for compositional and beta-diversity analyses [54] and principal coordinate analysis (PCoA) was used to visualize community differences in ordination space. We used 16S rRNA gene copy numbers, which can vary in number between the chromosomes of different bacterial and archaeal species and higher taxonomies [55], of *Aphanizomenon*, *Dolichospermum*, and *Microcystis* spp. from the University of Michigan ribosomal RNA database (rrnDB) [56] to infer putative cyanoHAB species cellular abundances (cells mL^-1^; more information in Supp. Methods) [57].

These data were used to bin samples into high and low cyanoHAB abundance (high: >1,000 cells mL^-1^ *and* during bloom time period; low: all other samples).

Microbiome and metatranscriptome compositional differences between statistical groups were performed with PERMANOVA, multivariate homogeneity of group dispersions, and ANOSIM tests with the vegan package (v2.6-4)[12]. Betadisper and permutest functions in vegan yielded a significant *p*-value (*p* < 0.05) for the two groups binned by cyanoHAB abundance, indicating significantly different dispersion of microbial communities between the groups. Therefore, as has been previously recommended [58], ANOSIM was used to test for community compositional differences between groups of high and low cyanoHAB abundance in taxa of microbiomes and metatranscriptomes. PERMANOVA was used to test for significant microbiome and metatranscriptome differences between samples taken in each season (pre-bloom in May and during the bloom in August) and between the four environmental types (spring water, river, lake, and reservoir).

### mRNA-seq data bioinformatics and statistics

Demultiplexed metatranscriptome reads were uploaded to the CosmosID (Germantown, Maryland, U.S.A.) microbiome bioinformatics HUB (see Supp. Methods for full description of CosmosID taxonomic and functional annotation pipeline) for taxonomic and functional profiling and linear discriminant analysis (LDA) effect size (LEfSe) analysis [59]. Taxonomic annotations of transcripts using GTDB quantified gene expression by individual bacterioplankton taxa at the time of sampling. Transcript functions were annotated with enzyme commission, Gene Ontology (GO) terms, protein family (pfam) hidden Markov model-based, and MetaCyc metabolic reconstruction databases, which are all integrated in CosmosID. LEfSe analyses were performed in the CosmosID-HUB to identify transcripts, grouped by taxa (from taxonomic pipeline) and function, that were significantly related to cyanoHAB abundance. Reference genomes (those that mRNA mapped to; basis of taxonomic annotations) of strains that were positively correlated with cyanoHAB abundance were accessed in CosmosID and searched for thiamin-related genes (Supp. Methods).

Per-sample functions of interest (GO terms and thiamin-related Pfam annotations) were exported as normalized counts (copies per million; CPM) and manually curated to assign higher-level functional descriptions (Supp. Methods). To examine the up-and down-regulation of gene expression in high and low cyanoHAB abundance bins, functionally annotated transcript CPM were corrected to z-scores [60], which were averaged across samples in each cyanoHAB abundance bin. *p*-values were adjusted to false discovery rate q-values indicating differential gene expression (q < 0.05) from per-sample z-scores with the pnorm function in R (two-sided tests; Data Availability statement for code). Metatranscriptome-based community composition and beta-diversity plots were produced following the same methods as for 16S taxonomy with microViz. For statistical tests including redundancy analysis and Spearman correlations between gene expression and TRCs and cyanoHAB abundances, the functionally annotated CPM abundance matrix was CLR-transformed (see Supp. Methods for full analyses steps).

PERMANOVA, ANOSIM, betadisper, and permutest functions were used following the same steps as for 16S taxonomy to assess between-group differences in transcriptionally active bacterioplankton.

## Results

### Dissolved TRC concentrations, nutrients, and bacterioplankton abundance

Median concentrations of dissolved thiamin-related compounds (TRCs; thiamin, HET, cHET, AmMP, HMP, and bacimethrin; i.e., not particle or cell associated) varied by 1 to 2 orders of magnitude across all UKB sample sites, and concentrations of thiamin and thiazole congeners were 1-3 orders of magnitude greater than those of pyrimidine congeners, including bacimethrin (Figure 2A; Table S1). Mean and median picomolar (pM) concentrations of most TRCs in May (pre-bloom time period, or pre-bloom period) differed from those in August (bloom time period, or bloom period) and greatly varied by sampling environments, being lowest in spring water and highest in reservoir water for each TRC (Figure 2; Tables 1 and S1). Based on Wilcoxon signed-rank tests, mean concentrations did not significantly differ between the pre-bloom and bloom periods of any TRCs. Mean TRC concentrations of thiamin, pyrimidine congeners, and bacimethrin were highest during the bloom period, while mean concentrations of thiazole congeners were highest during the pre-bloom period. The high TRC interquartile ranges in rivers relative to other UKB environments was driven by the low TRC concentrations observed in the Wood and Williamson Rivers compared to the Sprague and Link Rivers (Figure 2A; Table S1). The Link River (UKL outflow) also showed much higher concentrations of pyrimidine TRCs, including bacimethrin and especially during the bloom period (Figure 2A; Table S1).

**Figure 2.**
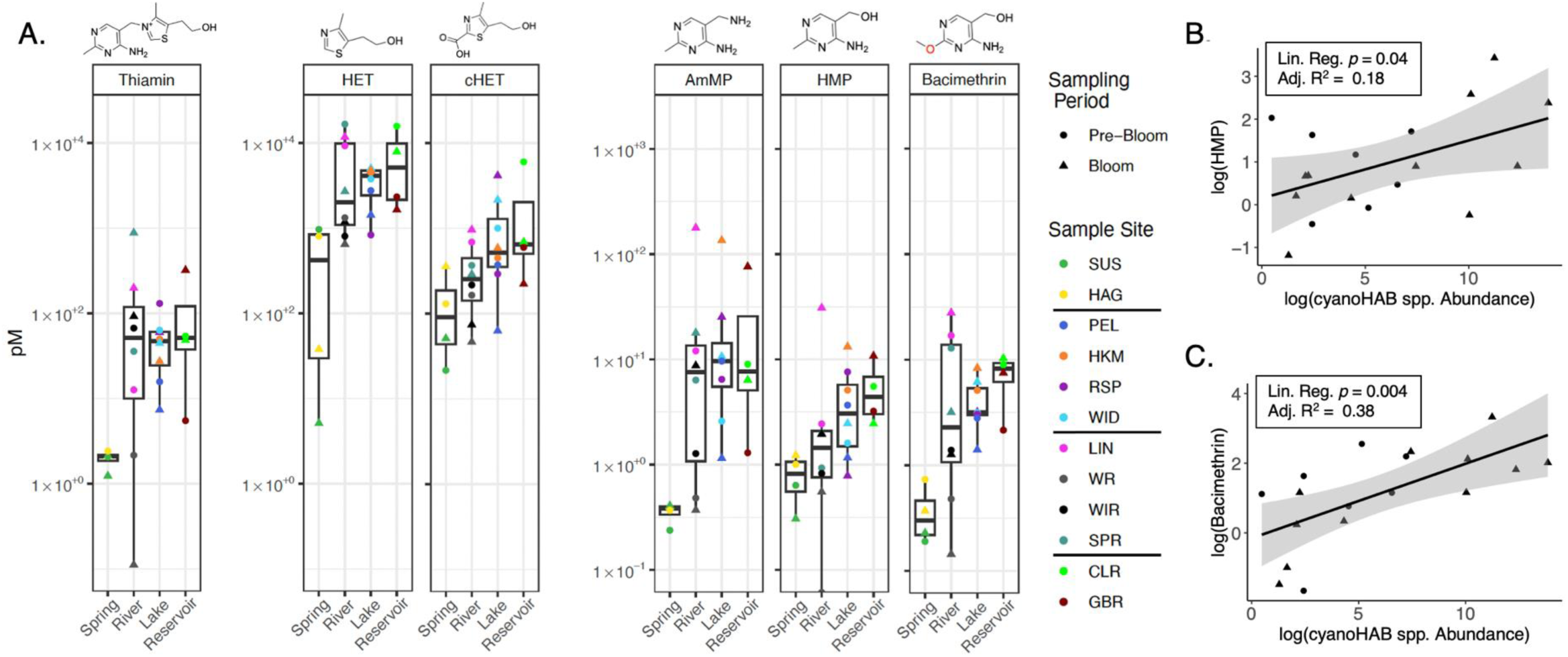
TRCs vary across sampling environments and correlate with the abundance of cyanoHAB spp. (A) TRC concentrations (picomolar; pM) presented as both discrete measurements and summary boxplots binned by sample environments. Lines are placed in between sample sites in color key to distinguish sampling environments (same order as shown in panel A x-axis). Chemical structures of TRCs are displayed above each compound’s boxplot. Scales differ between thiamin, thiazole, and pyrimidine TRCs to allow for comparability between individual sample points and median concentrations under the pyrimidine TRCs. Scatterplots displaying linear associations between log-transformed cyanoHAB spp. abundance (cells ml^-1^) and log-transformed (B) HMP and (C) bacimethrin concentrations. Linear regression results *(lm* in R function) are shown in boxes. In all panels, shapes correspond to sampling time periods: “Pre-Bloom” = May 2023; “Bloom” = August 2023.

Unlike TRCs, ratios of dissolved inorganic nitrogen (nitrate + nitrite + ammonium) to dissolved inorganic phosphorus (phosphate; N:P) differed greatly between pre-bloom and bloom periods (Figure S1). During the pre-bloom period, 8 out of 12 samples showed N:P values less than the Redfield ratio, suggesting N limitation (Figure S1) [61, 62]. During the bloom period, all samples except Clear Lake displayed N:P far above the canonical Redfield ratio of 16:1, suggesting P limitation on primary productivity, though typical N:P ratios in lacustrine environments are higher than those in the ocean [63, 64]. Further, during the bloom period, ammonium concentrations were high and phosphate concentrations were low (Figure S1) [65].

Bacterioplankton absolute abundances (total 16S gene copies mL^-1^ of sampled water) did not correlate with N:P concentration ratios (Spearman *p*-values > 0.05). Instead, bacterioplankton absolute abundances and TRCs each followed similar trends in magnitude across sampling environments, with increasing TRC concentrations and bacterioplankton absolute abundances from spring water to reservoirs (Figures 2 and S2). Bacterioplankton absolute abundances were also significantly and positively correlated with concentrations of all TRCs (all pairwise Pearson *p* < 0.05; all variables log-transformed). These results reflect a tighter coupling between bacterioplankton communities and TRC concentrations than between either of these factors and nutrients in UKB.

### CyanoHAB-related seasonal conditions and water quality

Cell abundances of cyanoHAB species (cyanoHAB abundance; cells mL^-1^) peaked in UKL, its outflow, and the reservoir sites during the bloom period (Figure S3A), and as with bacterioplankton absolute abundance, was positively correlated with bacimethrin concentrations (bolded brackets in Figure S3A). Linear regression results also showed that cyanoHAB abundance significantly predicted levels of bacimethrin and its non-toxic analog HMP (Figure 2B,C), indicating positive correlations between cyanoHABs, HMP, and bacimethrin. The most abundant cyanoHAB species were filamentous *Dolichospermum* spp. and *Aphanizomenon* NIES81. *Dolichospermum* spp. displayed cell abundances as high as 1×10^6^ cells mL^-1^ in the Gerber Reservoir (Figure S3A).

During the pre-bloom period when cyanoHAB abundance was low, USGS data indicated extensive hydrological mixing between UKL and the Williamson and Sprague River tributaries, based on high tributary discharge and lake surface elevation (Figure S4A,B) at this time. Outflow of UKL in the Link River displayed lower and more variable pH and DO concentrations during the pre-bloom period than the bloom period (Figure 1B,C), suggesting suppression of DO-generating primary productivity by elevated hydrological input from UKL tributaries. During the bloom period in August, cyanoHAB abundance was relatively high (Figure S3A), but the probability of poor water quality conditions was low based on surface elevation (∼1,261.7 m; Figure S4A) [46]. Link River pH and DO data suggested a small increase in primary production (high pH and DO) during August sampling, following the main bloom and subsequent crash that probably occurred between our sampling periods in late-July and early-August (Figure 1B,C) [46, 66, 67]. We also found no expression of cyanotoxin production genes in metatranscriptomics results, including *mcyE* [42] and any CosmosID functional annotation terms that included “microcystin” across all databases, despite high abundance of typical toxin-producers *M. aeruginosa* and *A. flos-aquae* (Figure S3A). These results suggest that the increase in cyanoHAB abundance during the bloom period did not degrade water quality.

### Connections between cyanoHAB abundance and bacterioplankton gene expression

When cyanoHAB abundance was high, gene expression was dominated by species of *Nostocaceae*, *Cyanobiaceae*, *Nanopelagicaceae*, *Microcystaceae*, and filamentous *Pseudanabaenaceae*. Conversely, when cyanoHAB abundance was low, gene expression was dominated by heterotrophic bacterial taxa such as *Burkholderiaceae* (two separate families: *Burkholderiaceae* and *Burkholderiaceae_B*), *Nanopelagicaceae*, and *Spirosomaceae* (Figure S6). This difference in metatranscriptomes was confirmed with ANOSIM (*p* = 0.003). However, 16S-based microbiome compositions did not significantly differ with changing cyanoHAB abundances (ANOSIM *p* > 0.05), despite visual separation of high and low cyanoHAB bins in ordinations of 16S-based microbiomes (Figure S3B,C). Microbiome compositions instead primarily differed by sampling environment and season (PERMANOVA *p*-values = 0.004 and 0.001, respectively). This statistical discrepancy suggests that, in contrast to the taxonomic compositions of metatranscriptomes (i.e., which taxa were actively transcribing genes), variability in DNA-based microbiome composition among habitats and seasons is greater than changes to composition driven by cyanoHABs (Figure S6).

Redundancy analysis results showed that cyanoHAB abundance, thiamin and pyrimidine concentrations (*p* < 0.05), and, to a lesser extent, bacimethrin concentrations (0.10 > *p*-value > 0.05; Figure 4B), predicted taxonomic compositions of metatranscriptomes. While its magnitude was low, the significance of the thiamin vector shows that thiamin concentrations influence taxonomic composition of transcriptionally active bacterioplankton. Other variables like sampling period (pre-bloom or bloom period), DO, and thiazole TRC concentrations displayed individual vector *p*-values >> 0.05, yet were significant to the overall prediction of metatranscriptome taxonomic compositions (model statistics in top-left box of Figure 4B).

Importantly, average NextSeq read quality and sequencing depth (∼30-80 million reads sample^-1^) were both high, indicating that differences between metatranscriptomes were driven by recognized environmental factors rather than concerns regarding sequencing fidelity (Figure S7).

LEfSe analysis identified bacterioplankton strains with gene expression levels that were enriched under high and low cyanoHAB abundance. Reference genomes for these taxa (i.e., those to which mRNA mapped; Supp. Methods) were queried for thiamin cycling genes (bolded genes in Figure 3A) to infer potential susceptibility to bacimethrin-mediated allelopathy (Figure 4A). Under high cyanoHAB bloom conditions all enriched cyanoHAB taxa, including *D. circinale*, *D. heterosporum* (annotated as *A. flos-aquae* on NCBI; Supp. Methods), and *M. aeruginosa*, were putative thiamin prototrophs (Table S2, Figure 4A), and thus likely resistant to bacimethrin. These putitively prototrophs were also among the most transcriptionally active taxa under high bacimethrin concentrations, suggesting a selective pressure against auxotrophic populations at these sites. Other abundant prototrophic taxa under high cyanoHAB conditions included *Pseudanabaenaceae* and picocyanobacteria *Cyanobium*, *Vulcanococcus*, and *Synechococcus* in Clear Lake, Link River, Rattlesnake Point, and Hank’s Marsh (*Cyanobiaceae* family; Figures 4A and S6, Table S2). Almost all reference genomes for these taxa contained the HMP biosynthesis gene *thiC* (Figure 3A; Table S2).

**Figure 3.**
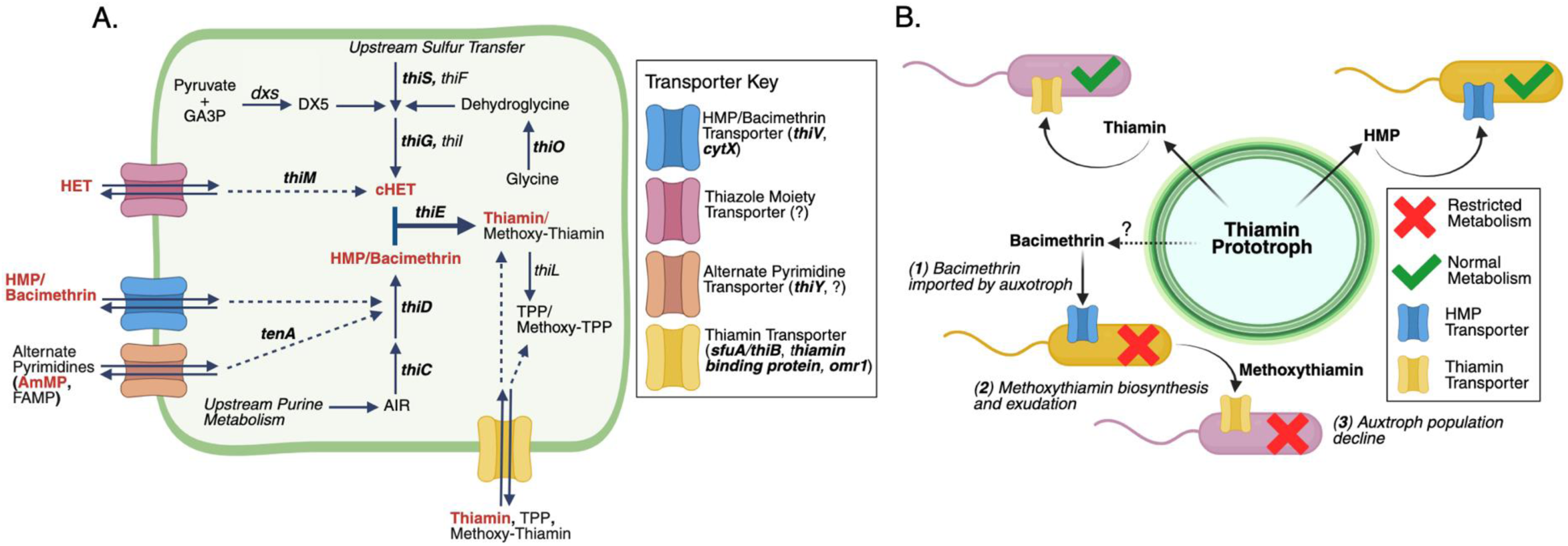
Routes of thiamin biosynthesis and salvage and the hypothesized impact of thiamin antagonists on UKB surface water bacterioplankton communities. (A) A generic cell (outlined in green) that contains all possible routes of thiamin biosynthesis (solid black arrows) and salvage (dashed black arrows). Red bolded labels indicate TRCs that were measured in this study and black bolded genes indicate those that were measured by hidden Markov models in reference genomes (see Supp. Methods for details). *“dxs*” = 1-deoxy-D-xylulose-5-phosphate synthase; “AIR” = 5-aminoimidazole ribonucleotide; “GA3P” = glyceraldehyde-3-phosphate; “TPP” = thiamin pyrophosphate. (B) A conceptual diagram showing the hypothetical metabolic impact of bacimethrin and methoxy-thiamin on thiamin auxotrophs. “?” indicates that the source of bacimethrin, whether from prototrophic bacterioplankton or an unidentified environmental source, is unknown, but would nevertheless impact bacterioplankton that import thiamin antagonists.

**Figure 4.**
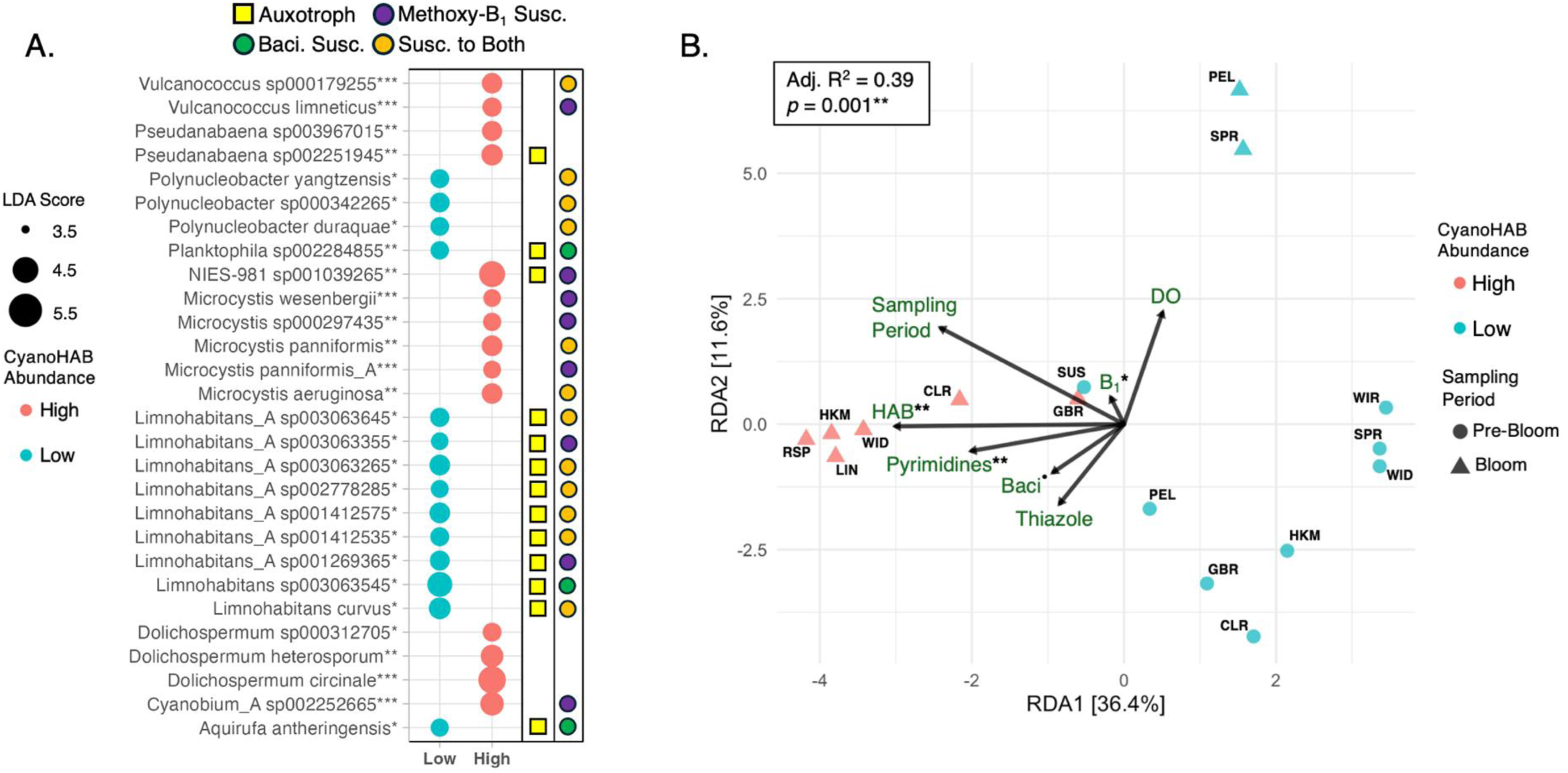
CyanoHABs and TRCs influence the gene expression of UKB bacterioplankton. (A) A bubble plot of LEfSe results that tested which strains (mRNA taxonomic annotations) most significantly differentiated high and low cyanoHAB abundance bins based on linear discriminant analysis (LDA) scores and their significance (*p* < 0.05*, *p* < 0.01**, *p* < 0.001***; attributed to each strain). Bubble size corresponds to the magnitude of LDA scores, where LDA magnitudes increase linearly with the degree of enrichment of each strain in its respective cyanoHAB abundance bin (or how predictive each strain is of that bin), though all taxa are significantly enriched in low or high statistical bins. Yellow squares show which strains are putative thiamin auxotrophs based on their reference genomes and the absence of a yellow square indicates thiamin prototrophs. Colored circles represent bacimethrin and methoxy-thiamin susceptibility based on the presence of HMP and thiamin transporters in reference genomes, respectively. (B) Constrained principal component analysis (RDA; also known as redundancy analysis) displaying differences (based on Aitchison distance) between strain-level taxonomic compositions of gene transcripts and explanatory variables that significantly (based on RDA model results in top-left box) predict taxonomic compositions of transcripts. Individual *p*-values of explanatory variables are shown next to each vector name (*p* < 0.1^•^*, p* < 0.05*, *p* < 0.01**).”HAB” = indicator variable (presence/absence) for high cyanoHAB abundance;“DO” = dissolved oxygen concentration,”Pyrimidines” = log-transformed HMP + AmMP concentrations, “Thiazole” = log-transformed cHET + HET concentrations, “Baci” = log-transformed bacimethrin concentrations, and “B_1_” = thiamin concentrations.

Conditions of low cyanoHAB and bacimethrin favored HMP auxotrophic strains of *Limnohabitans* spp. (*Burkholderiaceae* family), nearly all of which were linked to reference genomes that lacked *thiC*, but contained pyrimidine congener transporter genes *thiV*, *thiY*, and/or *cytX* [7] (Figure 4A, Table S2). Other taxa favored under low cyanoHAB conditions included *Polynucleobacteria* and Planktophilia spp. whose reference genomes showed varying levels of auxotrophy and susceptibility to bacimethrin (Figure 3B).

### CyanoHAB abundance and thiamin biosynthesis gene expression

Expression of thiamin biosynthesis genes was elevated when cyanoHAB abundance was high, including families of genes coding for thiazole and HMP biosynthesis (Figure 5A) and the thiamin genes, *thiC*, *thiF*, and *dsx* (Figure 5B). This pattern was particularly strong at Rattlesnake Point during the bloom period (“RSP_B” in Figure 5B; Table S3). This site also contained high relative abundances of *Cyanobiaceae*, *Pseudanabaenaceae*, *Microcystaceae* transcripts (Figure S4) and low concentrations of HMP and bacimethrin (Figure 2A), indicating a potential net depletion [68] of pyrimidine TRCs concomitant with thiamin biosynthesis and high non-filamentous cyanoHAB gene expression, as hypothesized for coastal marine environments. In Gerber Reservoir, a single strain of the filamentous cyanoHAB taxon, *D. circinale* represented >99% of the bacterioplankton gene expression during the bloom period (Figure S5). This site also displayed a significant upregulation in expression of biosynthesis genes for thiamin and other B vitamins (cobalamin and pantothenate–vitamins B_12_ and B_5_, respectively; Figure 5B, Table S3), implicating *D. circinale* as a major microbial source for B vitamins. Together, these data suggest that prototrophic cyanobacteria in UKB, including cyanoHAB species and non-HAB-forming cyanobacteria, produce the bulk of bacterioplankton-derived TRCs and are likely B vitamin sources for higher trophic levels in the basin.

**Figure 5.**
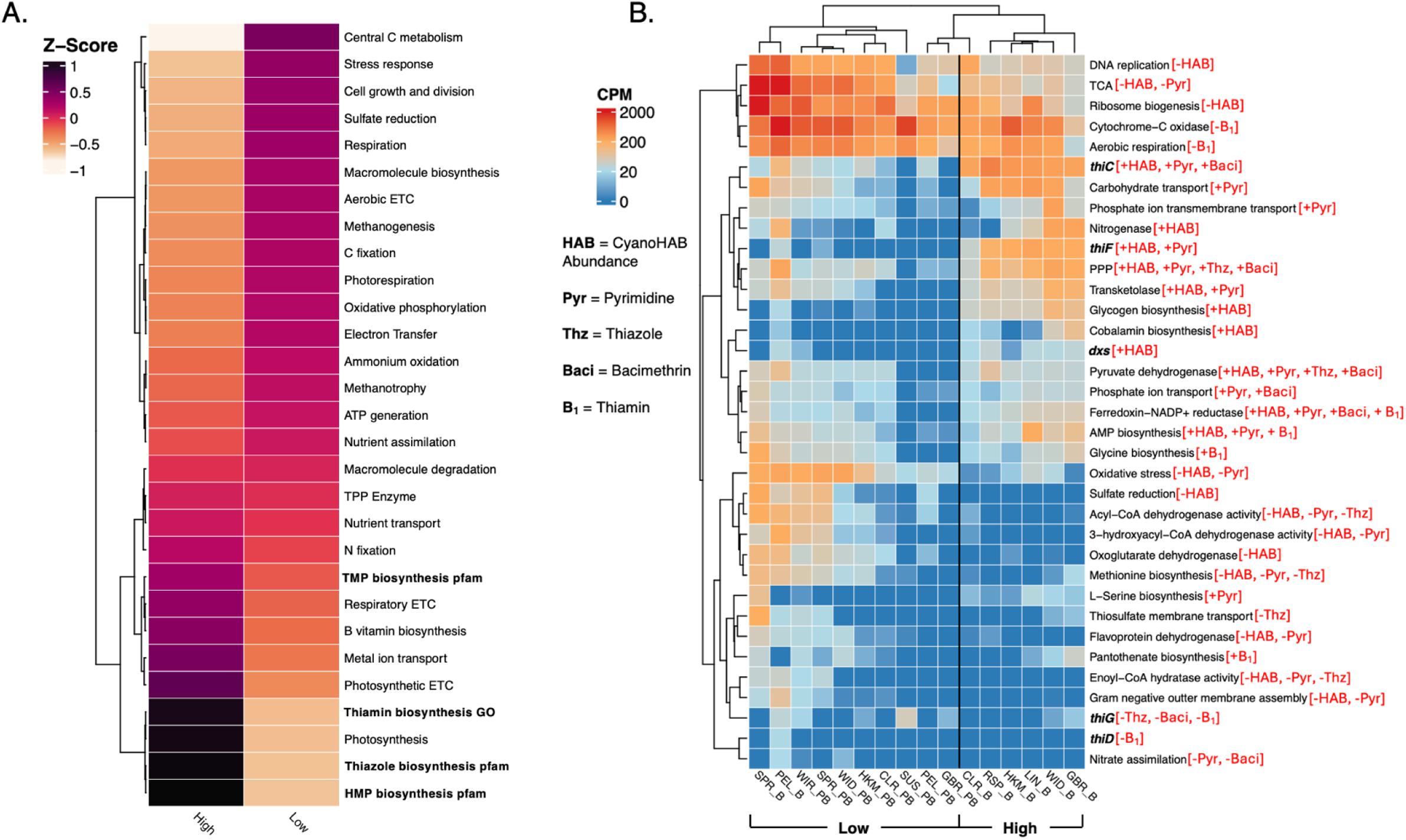
Bacterioplankton gene expression changes with differences in CyanoHAB abundance and influences concentrations of TRCs. (A) A heatmap of high-level functions (exact functional annotations displayed in panel B) and their associated z-scores averaged across all samples in each CyanoHAB biomass bin (high and low). Functions are ordered based on Euclidean distances of z-scores that were averaged across samples in each cyanoHAB abundance bin. (B) A heatmap of relative abundances (copies per million; CPM) for the top 35 most highly significant (lowest adjusted *p*-values) transcripts that correlated with cyanoHAB spp. abundances (cells mL^-1^) and TRC concentrations. Samples and functional annotations are ordered based on Bray-Curtis dissimilarity. Thiamin biosynthesis genes are bolded and were annotated with the pfam database; all other transcripts were annotated with the GO terms database. Significant correlations and the direction of correlations are shown in red bracketed text (see label key). The calculation of correlations (Spearman correlations between Aitchison distances of functional annotations and variables shown in figure key; see Supp. Methods) were taken from the Figure S12 correlogram. “PB” = pre-bloom and “B” = bloom. “PPP” = pentose phosphate pathway; “TCA” = tricarboxylic acid cycle; “Pyr” = correlated with at least one pyrimidine TRC: HMP and AmMP; “Thz” = correlated with at least one thiazole TRC: cHET and HET.

Expression of bacimethrin biosynthesis genes was not detected in our metatranscriptomes, possibly due to incomplete or ambiguous annotations. Many candidate genes, such as thymidylate synthase, present in both *Clostridium botulinum* and *Microcystis* spp. [20, 27], have broad biochemical functions, such as DNA synthesis [69], limiting their utility as diagnostic markers. Thiaminase I, found in *C. botulinum* but not *Microcystis* spp., was absent from all queried reference genomes. Other putative bacimethrin biosynthesis genes [20, 27] were similarly non-specific or absent, precluding confident detection of bacimethrin production via metatranscriptomics. Despite this, bacimethrin concentrations were positively correlated with cyanoHAB abundance (Figures 2C, S3A, 4B), strongly suggesting a microbial origin, potentially from prototrophic taxa active under high-cyanoHAB conditions.

### Correlations between patterns of gene expression and cyanoHAB-related TRC availability

Distinct gene expression patterns under high and low cyanoHAB abundances indicate that cyanoHABs substantially influence the metabolic activity of UKB bacterioplankton. Under high cyanoHAB abundance, there was elevated transcription of photosynthetic light reactions, the pentose phosphate pathway, the rubisco shunt, glycogen biosynthesis, gluconeogenesis, carbohydrate transporters, and thiamin biosynthesis (Figures 5A, 5B, S10, S12; see table S4 for how gene functions were categorized). These pathways reflect elevated photoautotrophic activity and *de novo* thiamin production. In contrast, low cyanoHAB conditions featured enhanced transcription of genes for aerobic respiration, central carbon metabolism (TCA, glyoxylate, and Calvin cycles), ammonium oxidation, and a range of heterotrophic processes including methanogenesis, methanotrophy, and assimilatory sulfate reduction (Figures 5, S9– S13; Tables S2, S3). Genes associated with cellular growth and division – such as ribosome, membrane, cell wall, and DNA replication pathways – also showed elevated expression in low cyanoHAB samples. The expression of fatty acid β-oxidation genes (e.g., acyl-CoA dehydrogenase) were negatively correlated with TRC concentrations and elevated under low cyanoHAB conditions (Figures 5B and S11). Oxidative stress gene expression was also highest in low-cyanoHAB samples and negatively correlated with cyanoHAB abundance and AmMP (Figures 5B, S11), suggesting redox stress may constrain thiamin biosynthesis or availability.

Several thiamin-dependent gene pathways also varied with cyanoHAB conditions.

Nitrogen fixation genes, which can be thiamin-dependent in cyanobacteria [70], showed only a modest increase under high cyanoHAB abundance (Figure 5A) despite strong expression of diazotrophic genes by filamentous and diazotrophic cyanoHAB taxa (Figure 4A). While nitrogenase transcription was positively correlated with cyanoHAB abundance (Figure 5B), it did not correlate with TRC concentrations, suggesting a limited link between nitrogen fixation and TRC dynamics. In contrast, expression of genes related to thiamin-requiring enzymes such as pyruvate dehydrogenase (TCA cycle) and transketolase (pentose phosphate pathway) positively correlated with cyanoHAB abundances and pyrimidine TRCs (Figure 5B). Thiamin-dependent 1-deoxy-D-xylulose-5-phosphate synthase (*dxs*; Figure 3A), which also supports cHET biosynthesis (Figure 3A), was similarly enriched under high cyanoHAB conditions (Figures 5A and S11) [71]. It is important to note, however, that 1-deoxy-D-xylulose-5-phosphate is also a precursor for isoprenoids and pyridoxal biosynthesis, in addition to cHET, yet regardless of the end product, *dxs* requires thiamin to function in the forward direction [72].

The expression of individual genes involved in HMP and thiazole biosynthesis showed opposing correlations with TRC concentrations, indicating that UKB bacterioplankton communities may uniquely cycle thiamin precursors depending on environmental conditions (Figure 5B red brackets; Spearman correlations in Figure S11). For example, *thiC* (HMP biosynthesis) and *thiF* (cHET biosynthesis) were positively correlated with TRC concentrations while *thiG* (cHET biosynthesis pathway) and *thiD* (HMP biosynthesis or salvage) were negatively correlated (Figures 3A and 5B). Notably, *thiG* was far more prevalent across cyanoHAB-associated reference genomes than *thiC*, suggesting a potential dominance of pyrimidine auxotrophy over thiazole auxotrophy in UKB bacterioplankton, consistent with findings from other aquatic systems [7] and the lowered average thiazole concentrations during the bloom period observed in UKB (Table 1). Additionally, *thiG* was highly expressed in the Sucker Springs groundwater site (Figure 5B), where expression of other thiamin biosynthesis genes was absent, implying that thiazole biosynthesis may be a more conserved trait across diverse UKB bacterioplankton, potentially explaining its weak correlation with cyanoHAB abundance.

**Table 1.** TRC concentration differences between the pre-bloom and bloom periods.

| TRC | Pre-bloom med. | Pre-bloom mean | Bloom med. | Bloom mean |
| --- | --- | --- | --- | --- |
| Thiamin | 25.7 | **36.7** ± 38.7 | 46.6 | **140.2** ± 251.9 |
| HET | 2538.0 | **4857.4** ± 5839.3 | 2169.5 | **3478.5** ± 3545.8 |
| cHET | 369.4 | **858.3** ± 1643 | 322.9 | **802.3** ± 1207.4 |
| AmMP | 4.5 | **5.0** ± 4.4 | 9.7 | **38.4** ± 59.7 |
| HMP | 2.0 | **2.7** ± 2.4 | 2.0 | **5.6** ± 9 |
| Bacimethrin | 2.9 | **4.8** ± 5.4 | 3.2 | **5.8** ± 7.8 |

*Standard deviations (sample, not population; calculated in Excel) are displayed after bolded mean values; “med.” = median.*

### A hypothetical model relating thiamin usage with hypereutrophic biogeochemistry

We developed a conceptual model (Figure 6) based on metatranscriptomic data to illustrate how bacterioplankton use TRCs under different cyanoHAB conditions. Gene pathways whose expression was correlated with cyanoHAB abundance, TRC concentrations including bacimethrin (see red brackets in Figure 5B), and putative allelopathic effects were integrated into cellular models representing the metabolic profiles of bacterioplankton under conditions of high and low cyanoHAB impact (Figure 6). While photosynthesis genes were expressed at all sites (Figure S13), differences emerged in ATP-expending pathways: photorespiration was upregulated under low cyanoHAB conditions, particularly at Pelican Bay (Table S3), where high groundwater input tends to buffer against cyanoHAB-driven water quality degradation (49, 81).

**Figure 6.**
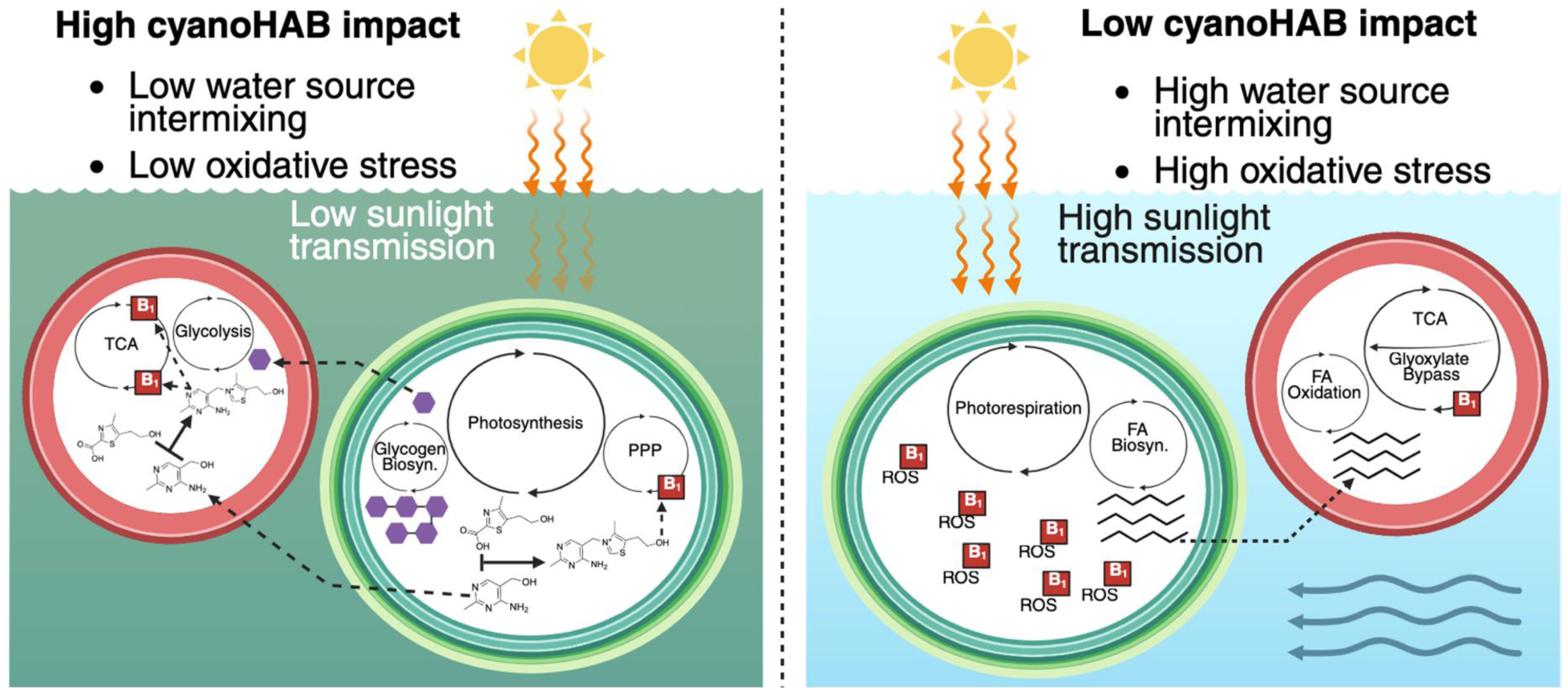
The impact of cyanoHABs correlates with cellular biochemical changes of bacterioplankton that influence how thiamin is used by cellular enzymes. The left and right sides of the figure display biochemical patterns of bacterioplankton gene expression in generic UKL habitats that, aside from the association between thiamin and ROS, are enriched in either high or low cyanoHAB abundance bins based on CosmosID LEfSe or differential expression (based on z-scores) results. The two panels do not portray the presence/absence of pathways, but rather the relative enrichment of the expression of genes found in pathways that are enriched in each scenario. Thiamin biosynthesis is portrayed to be increased under high cyanoHAB impact by the ligation of cHET and HMP to thiamin by green and red cells. Red “B_1_” squares indicate metabolic pathways that contain thiamin-requiring enzymes or, in the case with reactive oxygen species (ROS), the use of thiamin to degrade ROS. These thiamin-dependent pathways can each be inhibited by bacimethrin. Green and red cells portray generic algae and heterotrophic bacterioplankton. Cellular pathways depict transcriptional enrichment in conditions of different cyanoHAB impact. The bottom right panel horizontal arrows representing water current indicate greater input and mixing of multiple water sources such as groundwater and tributary water mixing with lake water or reservoir water. PPP = pentose phosphate pathway; TCA = tricarboxylic acid cycle; FA = fatty acid.

Sites with low cyanoHAB impact also showed elevated expression of gene pathways involved in reactive oxygen species (ROS) mitigation (Figures 5 and 6), consistent with increased light stress and oxidative pressure in the absence of shading by cyanoHAB surface scums (82–84). One such site, Pelican Bay, typically experiences elevated water quality due the high groundwater input in this area of the lake, which could have influenced thiamin-related patterns of bacterioplankton gene expression (Figure 6) [41, 73]. Conversely, sites with high cyanoHAB abundance likely experienced reduced light penetration caused by the surface scums (85, 86), further influencing algal and microbial photosynthetic dynamics (Figure 6).

Collectively, these observations support the biochemical framework proposed in our model and highlight how thiamin metabolism, oxidative stress response, and light-driven energy allocation differ across cyanoHAB gradients. This synthesis highlights the broader ecological consequences of cyanoHABs for microbial metabolic networks and (micro)nutrient cycling in eutrophic systems.

## Discussion

The observation that bacimethrin was present in all samples, with the highest concentrations (ca. 5-25 pM) measured under conditions of high cyanoHAB abundance (Figure 2), suggests that bacimethrin allelopathy in UKB surface waters has its greatest impact in the presence of cyanoHABs. In addition, we observed that dissolved concentrations of bacimethrin and its non-toxic analog HMP were positively correlated with cyanoHAB species abundances and nearly equimolar across diverse environments and cyanoHAB-related seasons, further indicating that the distributions of these sister-compounds are controlled by the same mechanism (Figure 2). This finding is not surprising because the known mechanism of action for bacimethrin-based allelopathy is competitive inhibition of HMP binding transporters and enzymes. Thus, we speculate that microbial uptake of both compounds via the same high affinity HMP transporters (such as ThiV and CytX) is the specific process controlling observed dissolved concentrations of bacimethrin and HMP.

A study using cell cultures of the ubiquitous marine bacterium *Candidatus* Pelagibacter st. HTCC7211, a known HMP auxotroph, reported half saturation constants (Km) for the HMP transporter ThiV to be within the range of observed bacimethrin and HMP concentrations in UKB (Km, 9.5 pM to 1.2nM; Figure 2) [74]. The similarity between the HTCC7211 Km and substrate concentrations found in the UKB provide strong evidence that the lower bounds of uptake affinity were governing dissolved concentrations in the environment. While previous research [74] has shown that HMP transporters are highly specific and not subject to competitive inhibition, bacimethrin was not evaluated and no HMP uptake kinetics studies have been conducted on freshwater microbial species. Further, we observed that high affinity HMP transporters were present across reference genomes (Table S2), especially in taxa (e.g. *Limnohabitans* spp.) whose transcriptional activity was negatively correlated with cyanoHAB abundance (Figure 4A).

The low observed concentrations of AmMP, HMP, and bacimethrin relative to thiamin and thiazole TRCs (Figure 2A) may reflect high cellular requirements for, and thus rapid uptake of, pyrimidine congeners by HMP auxotrophs. These low concentrations were observed in parallel with high transcriptional activity of the HMP biosynthesis gene *thiC* (Figure 5B); this could indicate that recently synthesized pyrimidine TRCs are more rapidly removed from the dissolved pool than other less essential TRCs, which would accumulate to higher concentrations without such rapid removal. This high microbial demand for pyrimidine congeners is supported by past research [7, 11] and would increase the allelopathic impact of bacimethrin (Figure 3A). CyanoHAB species were shown to have a particularly strong influence on pyrimidine (bacimethrin, HMP, and AmMP) TRC cycling, evidenced by linear regression, correlational, and gene expression results that showed increasing HMP and bacimethrin concentrations and *thiC* expression with high cyanoHAB abundance (Figures 2B,C and 5B). Additionally, we observed high transcriptional activity of putatively bacimethrin-producing *Microcystis* spp. indicating that this organism was present and highly transcriptionally active at the time of sampling in the UKB (Figures 4 and S6) [27].

The recent discovery that *Microcystis* spp. can produce bacimethrin in culture [27] suggests these cyanoHAB species may be important bacimethrin producers in UKB, especially given the high diversity of transcriptionally active *Microcystis* spp. (*M. aeruginosa*, *M. panniformis*, *M. wesenbergii*, and other unclassified species) that were enriched during cyanoHAB events (Figures 4A and S). *Microcystis* spp. use many strategies to rapidly respond to changing environmental conditions and outcompete other bacterioplankton and algae, which is exemplified by their expansive pan-genome [29] – their genetic repertoire of survival strategies – including microcystin production, high-affinity nutrient transporters, and efficient buoyancy mechanisms [75, 76]. *Microcystis* spp. reference genomes, which represented transcriptionally active strains in UKB, also displayed the highest variety of thiamin biosynthesis and salvage genes across all queried genomes (Table S2), indicating the potential for both full *de novo* thiamin biosynthesis and salvage of thiamin and pyrimidine TRCs (Figure 3A). Considering the adaptability of these taxa to the environment, the regulation of *Microcystis spp.* thiamin-cycling genes could be controlled by thiamin-dependent riboswitches [77], potentially allowing *Microcystis* spp. to switch between biosynthesis and salvage based on dissolved TRC availability. The ability of transcriptionally-regulated *de novo* thiamin biosynthesis could also enable *Microcystis* spp. to avoid importing toxic bacimethrin. Bacimethrin production may be another mechanism used by *Microcystis* spp., and potentially other cyanoHAB species, to dominate eutrophic watersheds.

Our data indicated that the influence of TRCs on bacterioplankton gene expression depends on the relative abundances of active thiamin prototrophs and auxotrophs in each sample. Although both thiazole and pyrimidine TRCs are essential and can be salvaged by cells, pyrimidines showed the strongest relationship with active community composition in this study (Figure 4B). This supports the idea that the specific type of thiamin auxotrophy – determined by the presence or absence of biosynthesis and salvage genes in reference genomes (Figure 3A) – influences which TRCs are most essential to a given microbial community. Our RDA results further support this, showing that pyrimidine TRCs and thiamin had a stronger influence on bacterioplankton gene expression than thiazole TRCs, which had no significant vectors in individual samples (Figure 4B). This may reflect the near absence of thiazole auxotrophs among reference genomes, all of which showed widespread capacity for thiazole biosynthesis and lacked the salvage gene, *thiM* (Figure 3A) including those correlated with cyanoHABs (Table S2). Our results suggest that thiazole TRCs are less essential and thus exert weaker influence on community structure than those associated with pyrimidine. This underscores the likely importance of bacimethrin allelopathy in the UKB system, as it targets HMP, the most limited TRC micronutrient identified in this study (Figure 2A and Table 1).

UKL environmental conditions reflect how different lake habitats, influenced by unique seasonal hydrological factors (groundwater and tributary flow) and cyanoHABs [41], foster bacterioplankton communities that cycle TRCs. For example, all pre-bloom UKL habitats displayed low thiamin biosynthesis gene expression (Figure 5A; Table S3) and high transcriptional activities of HMP auxotrophs (Figure S6; Table S2) that would require a continuous source of exogenous HMP. Conversely, as the summer progressed (Figures 1B,C), we hypothesize that autochthonous TRC production by cyanobacteria in UKL increased, concurrent with a lower UKL surface elevation and tributary input that reduces dispersal of cells and TRCs into the lake (Figure S4A,B). These observations from the UKB may parallel observations in global marine environments where diverse environmental conditions select for unique microbial communities that establish TRC concentrations in the dissolved pool [13, 68, 78]. In further support, dissolved TRCs across river habitats in the Sacramento River watershed, a system controlled by environmental conditions distinct from UKB, showed far lower thiamin concentrations (a median of 0.23 pM, compared to a UKB median across both seasons of 40.2 pM) and relative abundances of cyanobacteria than UKB [18]. Similar to other aquatic environments adjacent to urban centers [79], the unique environmental conditions within UKB habitats influenced the composition of bacterioplankton communities (Figure S3B). This environmental selection may have also had a measurable influence on TRC concentrations and the potential for bacimethrin allelopathy, based on which bacterioplankton populations were favored by specific environmental conditions of each UKB habitat and the thiamin-cycling genes encoded in the genomes of these populations.

The high bacimethrin levels concurrent with cyanoHABs in UKB negatively correlated with auxotrophic and globally ubiquitous heterotrophic bacterioplankton, suggesting that high bacimethrin and cyanoHAB abundance favors prototrophs and selects against auxotrophs (Figures 3B and 4A; Table S2). Our data showed a near complete depletion of the gene expression of auxotrophic *Limnohabitans* spp., during periods of increased cyanoHAB featuring prototrophic cyanobacteria species (Figures 4 and S6; Table S2). *Limnohabitans* (*Burkholderiaceae*) are globally ubiquitous in freshwater ecosystems and are capable of heterotrophic growth on organic exudates released by algae [80–82]. Reference genome analysis suggests that *Limnohabitans* spp. lack the *thiC* gene (Table S2), implying a dependency on exogenous pyrimidine TRCs, which could be alleviated by pyrimidine importation via ThiV, ThiY, and CytX transporters and the TenA salvage pathway (Table S2) [16, 78, 83]. While these strains could possibly avoid bacimethrin import by relying on the TenA salvage route to import exogenous AmMP (Figures 2A and 3A), their lack of transcriptional activity and reduced relative abundance at several sites dominated by cyanoHAB species (Figure S8) points towards an allelopathic mechanism inhibiting their metabolic activity (Figure 3B). This likely explains why the previously observed co-occurrence of *Limnohabitans* spp. with other types of algae [81] was not observed in UKB habitats.

The expression of other biogeochemically relevant genes related to C-cycling, energy acquisition, and cell stress across sites with varying cyanoHAB impact revealed additional metabolic pathways potentially influenced by TRCs and cyanoHAB abundances. These pathways included genes for fatty acid degradation enzymes, the glyoxylate cycle (or “bypass”), and oxidative stress response pathways (Figures 5, 6, and S11), all of which are connected to cellular thiamin demand. For example, elevated expression of fatty acid degradation genes in low cyanoHAB samples may reduce reliance on the thiamin-dependent pyruvate dehydrogenase, as fatty acids can directly enter the TCA cycle (Figure 6) [84]. Similarly, the glyoxylate bypass circumvents the need for thiamin-requiring 2-oxoglutarate dehydrogenase (Figure 6), whose associated gene expression was negatively correlated with cyanoHAB abundance (Figure S11). Periods of intensified oxidative stress experienced by UKB bacterioplankton may further deplete intracellular thiamin due to the thiazole ring’s role in deactivating ROS [85] (Figure 6).

Enrichment of photorespiration gene expression in low cyanoHAB sites provides additional evidence of oxidative stress and possible thiamin limitation (Figure 6). This energy-consuming process is typically associated with oxygen-rich conditions and has been proposed as a protective mechanism against photooxidative damage in cyanobacterial mats [86, 87]. Our observed negative correlation between photorespiration gene expression and dissolved thiamin concentrations (Figure S11), along with increased expression of these genes under low cyanoHAB conditions (Figure 5B and Table S2), support a link between photorespiration, cell stress, and limited environmental thiamin availability in these sites (Figure 6).

Using multiple methods, we find that active UKB bacterioplankton communities alter TRC availability and are uniquely susceptible to an allelopathic thiamin antagonist. Each thiamin-requiring metabolic pathway (see Figure 6) actively expressed by UKB bacterioplankton has the potential to be competitively inhibited by bacimethrin. As a result, changes in the expression of these pathways resulting from bacimethrin allelopathy exert strong biogeochemical influences on the persistence of cyanoHABs in UKB.

## Conclusion

CyanoHABs are a nuisance in freshwater ecosystems across the planet [88]. Our results point to the previously unrecognized importance of a toxic HMP analog, bacimethrin, as an allelopathic factor promoting cyanoHAB dominance and persistence. Microbial HMP auxotrophy is common, so while our results are specific to UKB, we expect that similar ecosystem-scale impacts occur in other watersheds similarly plagued by bloom-forming *Microcystis*, *Dolichosperum*, and *Aphanizomenon*. Bacimethrin allelopathy could be yet another mechanism for niche expansion exploited by cyanoHAB species, allowing these taxa to gain and maintain dominance in eutrophic watersheds.

## Acknowledgements

We thank the USFWS Klamath Falls Office including Rodger Gwiazdowski, Charlee Cramer, Ronald Twibell, and Christinia Kruse and U.S. Bureau of Reclamation staff Brock Phillips for their assistance with field sampling and project development. We thank Stephen Giovannoni for his support, mentorship, and use of his laboratory space. We thank Jeff Morré and Jaewoo Choi for assistance with mass spectrometry. Finally, we thank the broader community of Salmonid Thiamin Deficiency Complex scientists for helpful and inspirational conversations that have shaped the direction of this research (workshops funded by NCEAS Morpho, USGS John Wesley Powell Center for Analysis and Synthesis, and NSF). We thank Beth Ahner for her advice, support, and collaboration surrounding bacimethrin. This work was funded by United States Fish and Wildlife Service grant F22AC01810-01 and California Department of Fish and Wildlife grant Q2196012, both to Christopher P. Suffridge. Additional personnel funding for Christopher P. Suffridge was provided by National Science Foundation grant DEB-1639033.

Mass spectrometry instrumentation at the OSU Mass Spectrometry Center was supported by National Institutes of Health grant 1S10RR022589-1. The authors declare no conflict of interest.

## Data Availability

All microbial sequence processing, raw LCMS data, and data analysis scripts can be found here: https://github.com/ksmicrobe/UKB-Thiamin/tree/main/Manuscript_Code and all sequencing data (MiSeq and NextSeq) can be accessed with the following NCBI SRA BioProject ID: PRJNA1216032.

