## Supplemental methods and figures for "The allelopathic vitamin B1 antagonist bacimethrin impacts microbial gene expression in a hypereutrophic watershed dominated by cyanobacterial blooms"

Short Title: Bacimethrin allelopathy in a hypereutrophic watershed

### Extended Methods

*Detailed sampling and sample site description*

Surface water samples were collected at a depth of ~0.5 m using 1 L amber HDPE bottles (Nalgene). Surface water samples in the rivers and reservoirs were taken at knee height by wading into the water and Upper Klamath Lake samples were taken from a boat. All samples were prefiltered (100 µm mesh filter) and kept in the same 1 L bottles on wet ice until further processing. Within the same day of sampling, peristaltic filtration at a rate of 30 mL min$^{-1}$ was used to collect cells and particles onto 0.22-µm Sterivex filters (PES membrane, Millipore, Burlington, MA, U.S.A.) and volumes of filtered water were recorded. Up to 1 L of cell-free filtrate (depending on filter clogging from particles and suspended sediment) was collected in acid-washed and methanol-rinsed amber HDPE bottles for thiamin congener and bacimethrin analyses, which were stored in a -20 °C freezer. Immediately following filtration, 1 mL of RNA*later* (Thermo Fisher Scientific) was pipetted directly into Sterivex casings, which were then sealed and incubated in the dark for 5 minutes. Following incubation, sealed Sterivex cartridges were flash frozen in liquid $N_2$ and immediately placed in a -80 °C freezer. Samples were transported back to Oregon State University (OSU) on dry ice and were immediately placed in -80 °C upon arrival until the commencement of lab analyses.

In both sampling timepoints, the sampling location at Wood River was slow, shallow, and meandering whereas all other rivers were wider, deeper, and, based on visual observation, had substantially higher flow rates: from slowest to fastest, Williamson, Sprague, and Link River. Sites within Upper Klamath Lake also ranged in ecological characteristics. Pelican Bay has high spring water inputs and is generally less impacted by CyanoHABs compared to other locations in the lake, Williamson Delta and Hank's Marsh sites are each located within wetland habitats, and the Rattlesnake Point site is the least sheltered and most exposed to wind-induced water mixing. The Sucker Springs site is located along the shoreline of Upper Klamath Lake where spring and lake water mix, with the sample itself being mostly spring water, and Hagelstein Pond is located adjacent to the lake and is entirely spring water. Finally, the Clear Lake and Gerber reservoirs are each headwater lakes of the Lost River watershed.

*USGS data analysis*
All data was plotted between 05/01/2023 – 09/01/23 with the dataRetrieval USGS R package (1). For Upper Klamath Lake surface elevation, measurements were averaged between three sites: Pelican Bay (site code = 11505800), Rattlesnake Point (site code = 11505900), and Klamath Falls (site code = 11507000) and corrected from ft. to m. Discharge rates for the Sprague (site code = 11501000) and Williamson (site code = 11502500) Rivers were averaged and corrected from cubic ft. $s^{-1}$ to cubic $m^{-1}$. Raw pH and DO values were used for the Link River site (site code = 421401121480900). Water surface elevation (parameter code = 72275), discharge rates (parameter code = 00060), pH (parameter code = 00400), and DO (parameter code = 00300) were retrieved with the "readNWISuv" command and measurements were plotted with ggplot2.

*Dissolved TRC LCMS analysis*
An LCMS method was utilized to simultaneously measure thiamin congener and bacimethrin concentrations. Analysis was conducted using an Applied Biosystems 4000 Q-Trap triple quadrupole mass spectrometer with an ESI interface coupled to a Shimadzu LC-20AD liquid chromatograph. Applied Biosystems Analyst and ABSciex Multiquant software were used for instrument operation and sample quantification. A Poroshell 120 PFP, 3 × 150 mm 2.7 µm HPLC column (Agilent) with a Poroshell 120 PFP, 2 × 5 mm, 2.7 µm guard column (Agilent) was used for chromatographic separations. Chromotography conditions, mass spectrometry settings, and compound specific information have been previously published (ref 28, ahner). Samples were indiscriminately randomized prior to analysis. To compensate for matrix effects, $^{13}C$-labeled thiamin was used as an internal standard. LCMS analysis was conducted at the Oregon State University Mass Spectrometry Core Facility.

*DNA/RNA Sterivex extraction and sterilization steps*
Only RNase-free microcentrifuge tubes were used for samples containing RNA. Lab gloves were also frequently sterilized with Obliterase and 70% ethanol when moving in and out of the biosafety hood. Prior to extractions, Sterivex filters were briefly thawed at room temperature and cartridges were cracked open using pliers. The thawed liquid inside cartridges (a mixture of RNA*later* and sample) was poured into UV-sterilized Petri dishes and Sterivex filters were cut into smaller strips with sterile medical scalpels and autoclaved tweezers in Petri dishes. Additionally, Petri dishes, scalpels, and tweezers were sterilized with RNaseZAP (Thermo Fisher Scientific) to prevent RNA degradation. DNA and RNA were extracted from cut up Sterivex filters using a ZymoBIOMICS (Irvine, California, U.S.A.) DNA/RNA Miniprep Kit (any sample type). Negative controls of nuclease-free water were extracted and underwent all pre-sequencing steps for DNA and RNA samples in parallel with true samples to assess contamination.

*PCR and Illumina MiSeq preparation steps*
PCR was performed with the following conditions: 2 min initial denaturation at 94°C, 30 cycles of denaturation at 94°C for 15 s, annealing at 50°C for 15 s, and extension at 72°C for 15 s, followed 10 min at 72°C for a final non-cycled hold. PCR was performed in duplicate on 96-well

plates, with each well containing 1 µl template DNA, 5 µl primers, 12.5 µl Platinum II master mix, and 6.5 µl PCR water. Amplicons from each double well were then pooled together and purified with a QIAquick PCR purification kit (Qiagen; Venlo, The Netherlands). Following purification, all samples were pooled to an equimolar concentration based on Qubit fluorometer values and sent to the OSU Center for Quantitative Life Sciences (CQLS) for Illumina (San Diego, California, U.S.A.) MiSeq 2x250 paired-end high throughput sequencing (HTS). Prior to sequencing, amplicon quality and proper sizes were checked with a BioAnalyzer by the CQLS.

*T. thermophilus extraction and 16S absolute abundance workflow*

Prior to the addition of cut-up Sterivex samples into lysis tubes, 22.4 ng of *Thermus thermophilus* strain HB8 (~1x10$^{7}$ genomes; estimated based on DNA concentration of the stock and the size of the HB8 genome), supplied by the ATCC (Manassas, Virginia, U.S.A.), was pipetted into lysis tubes filled with DNA/RNA Shield. The concentration of the *T. thermophilus* stock was based on the mean of the asymptote values of each Qubit fluorometer concentration of a serial dilution of the stock (where average concentrations normalized across: 1x, 5x, 10x, 50x, 100x, 200x, and 500x dilutions), multiplied by each dilution factor. To allow for more thorough cell lysis and freeing of cells from Stervix filter fragments, bead beating was preliminarily performed for 5 minutes, followed by the transfer of 400 µl sample to a nuclease free tube (reduced crowding of filters in the lysis tube), 5 more minutes of bead beating, and the transfer of the remaining sample from the lysis tube to the same nuclease free tube.

All *T. thermophilus*-related bioinformatics code can be found in the Data Availability Statement link. Both 16S reference databases yielded nearly identical total *T. thermophilus* read counts (based on read counts of ASVs annotated to the *Thermus* and *Meiothermus* genera; Silva = 201,785 and GTDB = 201,792 *T. thermophilus* reads), showing that the database used did not influence the taxonomic annotation of *Thermus thermophilus* to a degree that would impact the final absolute abundance estimations. Silva-based *T. thermophilus* per-sample read counts were exported into Excel to calculate total 16S copies mL$^{-1}$ of water (absolute abundance) in each sample in accordance with previous methods (2) (Excel table of calculations in Data Availability

(T. thermophilus genomes added)

Statement GitHub link) and based on the equation: [] x (non −

(9 : " .%&'()*+ &,-./ ;1675 )

T. thermophilus reads) x[ < ], which yields 16S rRNA gene copies mL$^{-1}$. *T.* =>)+-'

?*-.)' @,-%'('A

*thermophilus* reads are multiplied by 2 to account for the *T. thermophilus* strain HB8 16S rRNA gene copy number of 2. For Pearson correlations between thiamin congener and bacimethrin concentrations and absolute abundances, all variables were log-transformed and tested for normality with Shapiro tests ($p$-values > 0.05).

To correct per-sample ASV relative abundances to ASV absolute abundances, data table cells of each column (representing samples) of the ASV compositional abundance matrix was multiplied by that sample's absolute abundance estimate and a phyloseq object was generated containing the absolute abundance ASV table. Four individual phyloseq objects were then created for each unique CyanoHAB genus (*Aphanizomenon* MDT14a, *Aphanizomenon* NIES81, *Microcystis* PCC-7914, and *Dolichospermum* NIES41) and 16S copies were corrected to cell numbers by dividing all values in each respective ASV table (absolute abundance) by the 16S copies genome$^{-1}$ estimates of each genus, based on the University of Michigan ribosomal RNA database (rrnDB) (3). The Silva reference database (v138.1) (4) was used to assign taxonomies to species of CyanoHABs (*Aphanizomenon*, *Dolichospermum*, and *Microcystis*) and for all *T. thermophilus* absolute abundance steps due to our analysis yielding improved 16S-based genus- and species-level taxonomic resolutions for these CyanoHAB species with Silva compared to the GTDB. Copy numbers of all *Aphanizomenon*, *Microcystis*, and *Dolichospermum* species were 5, 5, and 2 in the rrnDB, respectively. Spearman correlations were then calculated between the persample total CyanoHAB biomasses (cells $mL^{-1}$) of the four unique genera and bacimethrin concentrations. Each phyloseq object containing only cyanoHAB species within each of the three genera was then merged into one and a heatmap of CyanoHAB biomasses was generated with microViz.

*CosmosID metatranscriptomics taxonomic and functional analysis steps*
Taxonomic results are available as filtered (meet CosmosID internal filtering threshold), highconfidence reads and total reads, and only filtered reads were used in all analyses. The CosmosID genome database (GenBook) relies on >30,000 phylogenomically-organized microbial genomes (bacteria, archaea, viruses, protists, and fungi) with taxonomies based on the GTDB. GenBook database curation is based on the collection of genomes with high completeness, low contamination, and high intra-species diversity and their assembly into a reference phylogenetic tree. The genomes are split into n-mers (biomarkers) of variable length, which are phylogenetically classified as being either shared or unique across genomes so that shared genomic biomarkers represent the tree backbone and unique biomarkers represent tree leaves. Taxonomic annotations within CosmosID are performed with the patented KEPLER pipeline following quality control steps to trim low-quality bases and adaptors with BBDuk (5) and quality visualization with MultiQC (6) (Figure S7). The pipeline consists of splitting reads into k-mers and finding exact matches between query k-mers and reference biomarkers in the GenBook reference tree. A list of nearest reference strains is then generated from these matches and a probabilistic Smith-Waterman edit distance-scoring based algorithm is used to assign reads to strain-level taxonomies. The reference genomes that mRNA reads aligned to were those queried for thiamin cycling genes (if they were significantly associated with cyanoHABs based on LEfSe results). An iterative Maximum Likelihood Estimation algorithm is then implemented to provide final normalized read abundance and relative abundance estimates.

For functional annotations, the CosmosID pipeline consisted of quality controlled reads being translated and searched against the UniRef90 protein sequence database, provided by UniProt (7). Read mapping was performed by CosmosID following the methods of Franzosa, McIver (8) and gene families were annotated to MetaCyc (9) pathways, Pfam (protein family) protein domains, Enzyme Commission (EC) Enzymes, and GO (gene ontology) Terms (subset into molecular functions, biological processes, and all GO Terms). Functional abundances were available as both copies per million (CPM) and relative abundances. Lefse analyses were performed in the CosmosID-HUB to test relative abundances of strains (from taxonomic pipeline) and functional annotations (EC, GO Terms, and Pfam annotations) that were significantly enriched with CyanoHAB impact (high and low biomass bins mentioned prior). For the manual curation of functional results, functional annotations were categorized into 11 total higher level processes: B vitamin biosynthesis, central C, nutrient acquisition, cell stress, cell growth and division, macromolecule biosynthesis, biogeochemical (general C and nutrient cycling), energy generation, electron transfer, TPP enzyme, and macromolecule degradation in order to gain a comprehensive perspective on microbial ecological processes in the system and how they relate to thiamin (TPP) usage and cycling (see Table S4 for full collection of annotations). These data were made into a phyloseq object for further analysis.

*Reference genome thiamin-related gene searches*
To assess the genomic potential for thiamin cycling of transcriptionally active taxa that were associated with CyanoHABs, reference genomes (from taxonomic pipeline; see above) of bacterioplankton strains that were significantly enriched with high or low CyanoHAB presence were compiled. The CosmosID reference database only provides pfam-based hidden Markov models (HMMs), so only a subset of thiamin cycling genes could be queried in metatranscriptomes. Further, the pipeline doesn't allow the user to associate functional annotations with taxonomic identities of the taxa that expressed the transcript. To address these shortcoming, we used the full diversity of thiamin-cycling HMMs (10) (available on GitHub; see Data Availability Statement) from all Interpro databases to search reference genomes. For each reference genome, both the GTDB- and NCBI-based taxonomic annotations were noted. NCBI taxonomic annotations are available for all GTDB ones and generally differ slightly due to the precise nature of how the GTDB assigns taxonomy based on the phylogenomic tree of marker genes. Each reference genome was then queried for thiamin cycling genes using the HMMs from Table S1 in Paerl, Sundh (10). All genome links and raw results can be found in Table S2. Decoy HMMs were used to test for false positive hits where appropriate (*thiY*). All HMM searches were performed with hmmer (v3.3.2) (11) on the command line with hmmsearch set to an E-value threshold of $1x10^{-10}$. Strains, represented by reference genomes, were then assigned as being susceptible to bacimethrin or methoxy-thiamin if they contained HMP transporter genes (*cytX* and/or *thiV*) or thiamin transporter genes (*sfuA*/*thiB*, thiamin binding protein, and/or *omr1*), respectively. Thiamin auxotrophs were defined as strains whose reference genomes lacked at least one of the core thiamin biosynthesis genes: *thiC*, *thiG*, and *thiE*.

For all reference genomes of *Aphanizomenon*, *Microcystis*, and *Dolichospermum*, both the *thiO* and *thiG* HMMs came up as hits to the same gene, though *thiG* hits had lower E-values (higher confidence annotations). To investigate this further, we copy-pasted each gene (translated) from *thiO*/*thiG* hits from all CyanoHAB reference genomes into the NCBI protein BLAST GUI and ran them against the non-redundant protein sequences reference database, with each respective CyanoHAB genus selected under the "Organism" tab. All protein sequences had hits to both *thiO* and *thiG* protein sequences for each respective genus and NCBI results showed that these genes had been merged: the first half of the protein sequence was annotated as *thiO* and the second half as *thiG*. Though it's unknown whether these genes could have been fused in each of the CyanoHAB chromosomes, interpreted this result as evidence for each species of CyanoHAB being a thiamin prototroph (containing the full thiazole biosynthesis pathway).

*RNA-based statistical analysis*
Redundancy analysis (RDA), which is a version of principal component analysis (PCA) where principal components are constrained to be linear combinations of explanatory variables (12), was performed with the vegan (v2.6-4) package (13). RDA was used to test the amount of RNAbased community compositional variance explained by the presence/absence of cyanoHABs, sampling time (prior to or during cyanoHAB bloom seasons), DO, the log-transformed sum of pyrimidine (HMP + AmMP) and thiazole (cHET + HET) congeners, and log-

transformed thiamin and bacimethrin concentrations. Chemically similar thiamin congeners were summed to reduce multicollinearity. ANOVA tests were used to test model significance and variance inflation factors were checked to all be below 10, indicating limited multicollinearity of variables. For the CPM-based functional heatmap, function and sample ordering (seriation) was performed within the microViz "comp_heatmap" command based on Bray-Curtis dissimilarities. A matrix of Spearman correlations between CLR-transformed functional annotations (CPM) and untransformed thiamin congeners, bacimethrin, and 16S-based absolute abundances was constructed with microViz, subset to only contain functions with at least one significant (adjusted $p < 0.05$) correlation. *p*-values were corrected with the Benjamini and Hochberg method to control for false-positive correlations.

**Supplemental Figures**

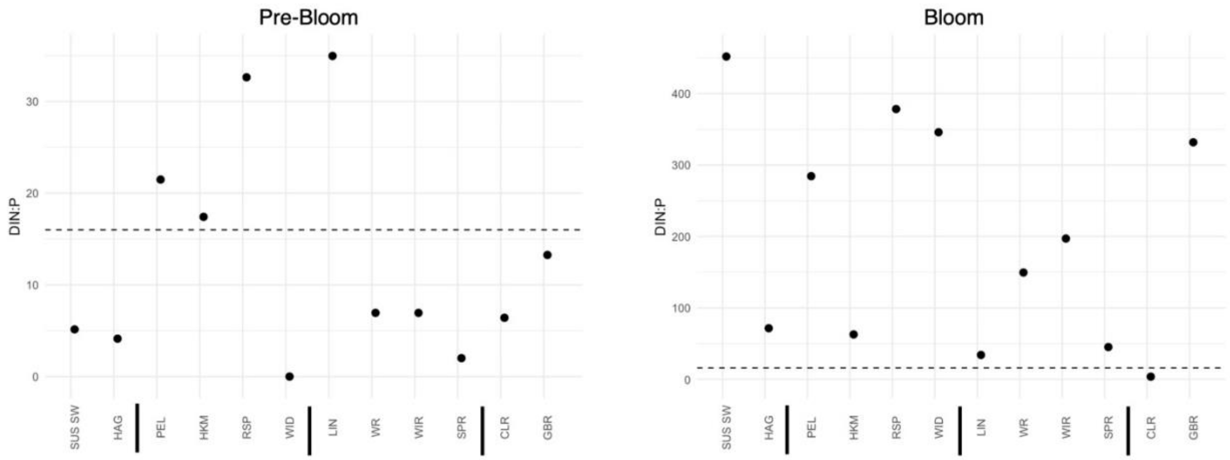


**Figure S1.** Nutrient ratios change based on the sampling season. µM nutrient ratios of DIN:phosphate (P) across sample sites in the pre-bloom and bloom time periods. Dashed lines are drawn at 16 µM to indicate the Redfield ratio of N:P of 16:1.

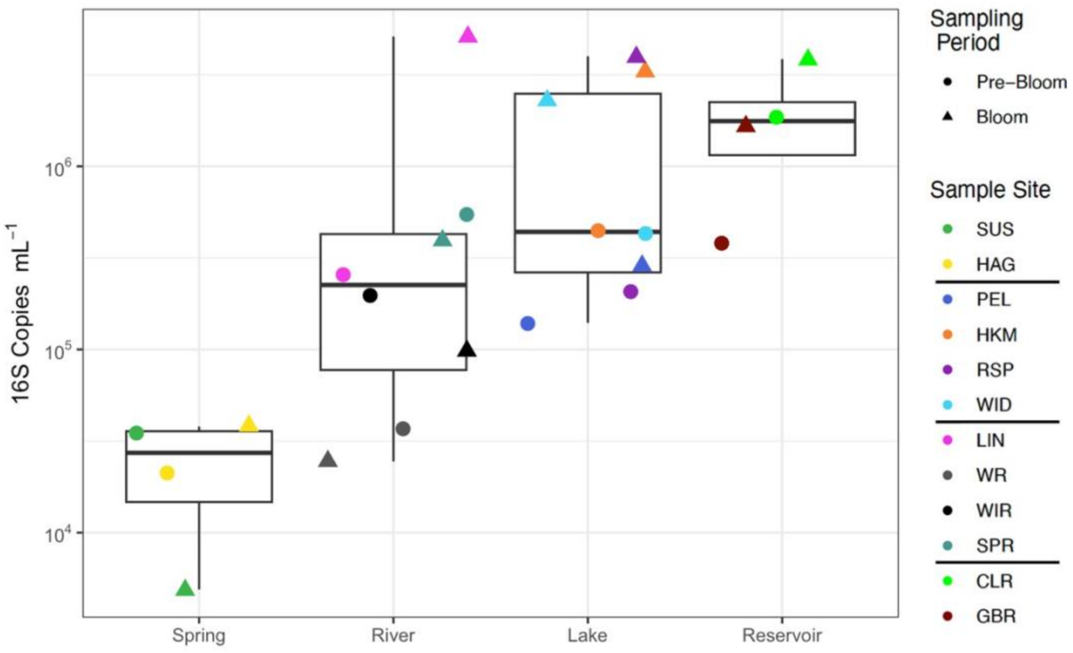


**Figure S2.** Absolute abundance of 16S copies mL$^{-1}$ based on *T. thermophilus* spike-in internal standard results . Shapes correspond to sampling time periods; “Pre-Bloom” = May 2023, “Bloom” = August 2023. Lines are placed in between sample sites in color key to distinguish sampling environments (same order as shown in panel A x-axis).

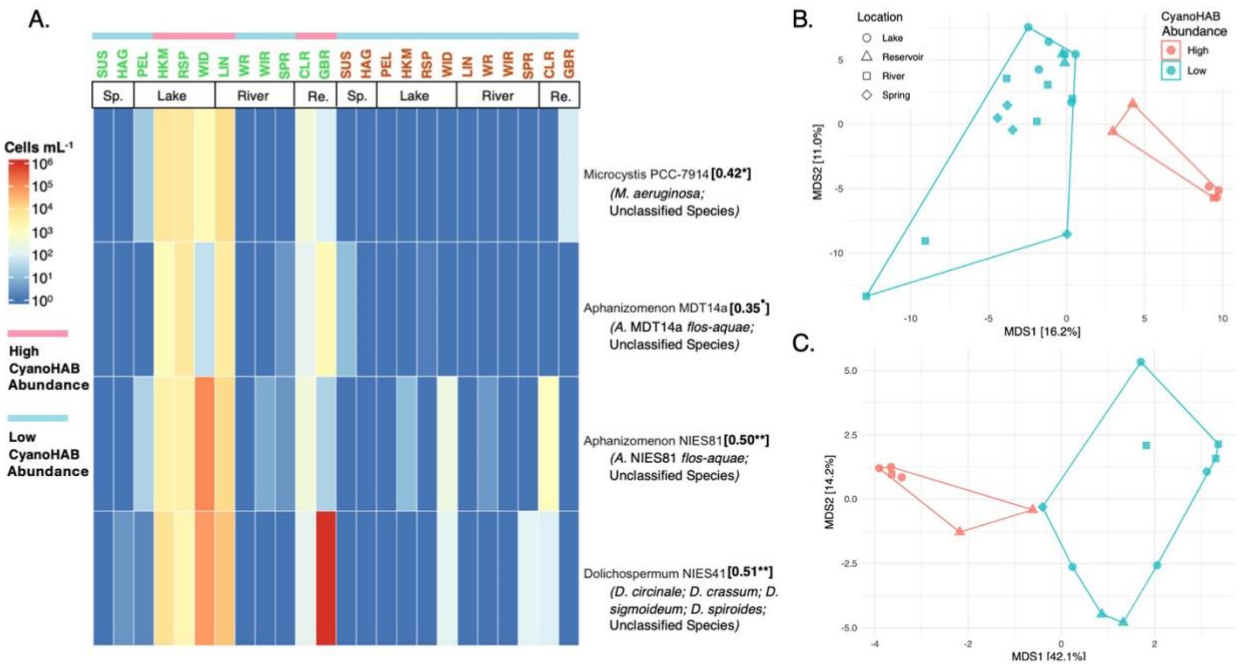


**Figure S3.** CyanoHAB cellular abundances (cells mL$^{-1}$; abundance) are highest during the bloom time period and influence bacterioplankton. (A) Heatmap showing the abundance of each unique cyanoHAB genus based on 16S gene absolute abundances and copy numbers (see Methods). Samples taken during the bloom period (colored green) are shown first followed by pre-bloom (colored orange) samples and cyanoHAB abundance bins are indicated above site names. Twosided Spearman correlations and significance levels *(p* < 0.1$^{\bullet}$, *p* < 0.05*, *p* < 0.01***)* are shown in bolded brackets next to each cyanoHAB genus and represent correlations between the total biomass of all ASVs in each of the four unique genera and bacimethrin concentrations. PCoA, or metric multidimensional scaling (MDS), ordinations were constructed to display differences (based on Aitchison distance) between (B) ASV-level 16S-based microbial community compositions and (C) strain-level taxonomic compositions of gene transcripts across sites. Shapes and colors represent sampling environments ("Location") and cyanoHAB abundance bins, respectively.

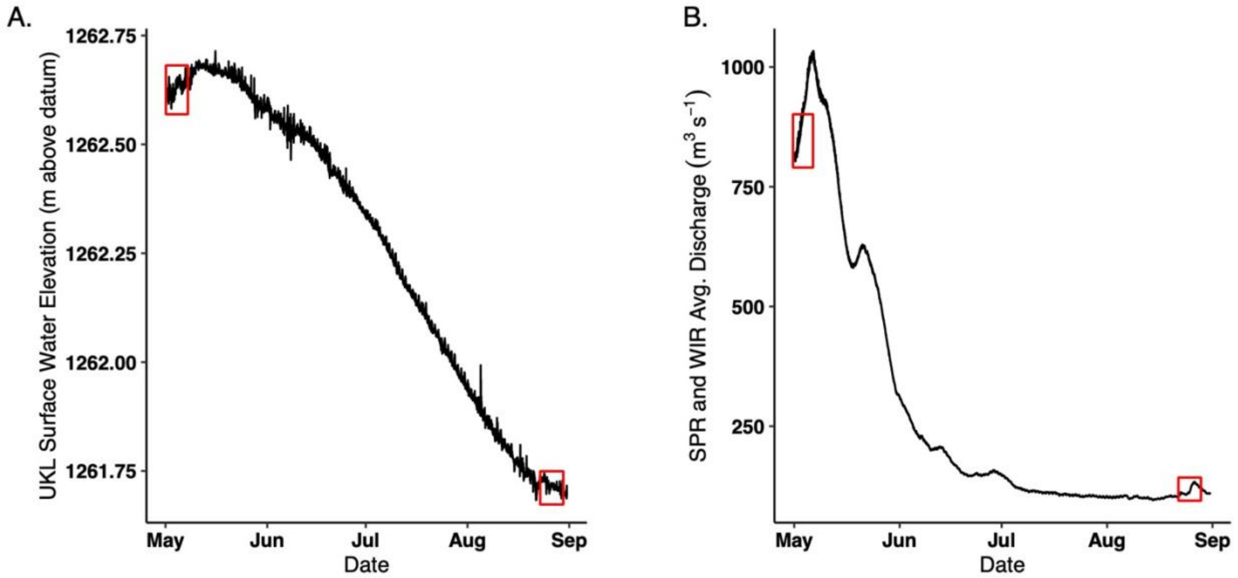


**Figure S4.** Water quality related factors change in UKL across the summer. (A) Surface water elevation and (B) tributary discharge (Sprague and Williamson Rivers) were plotted from USGS hourly water quality monitoring stations between May 5$^{th}$ and Sept. 1$^{st}$, 2025.

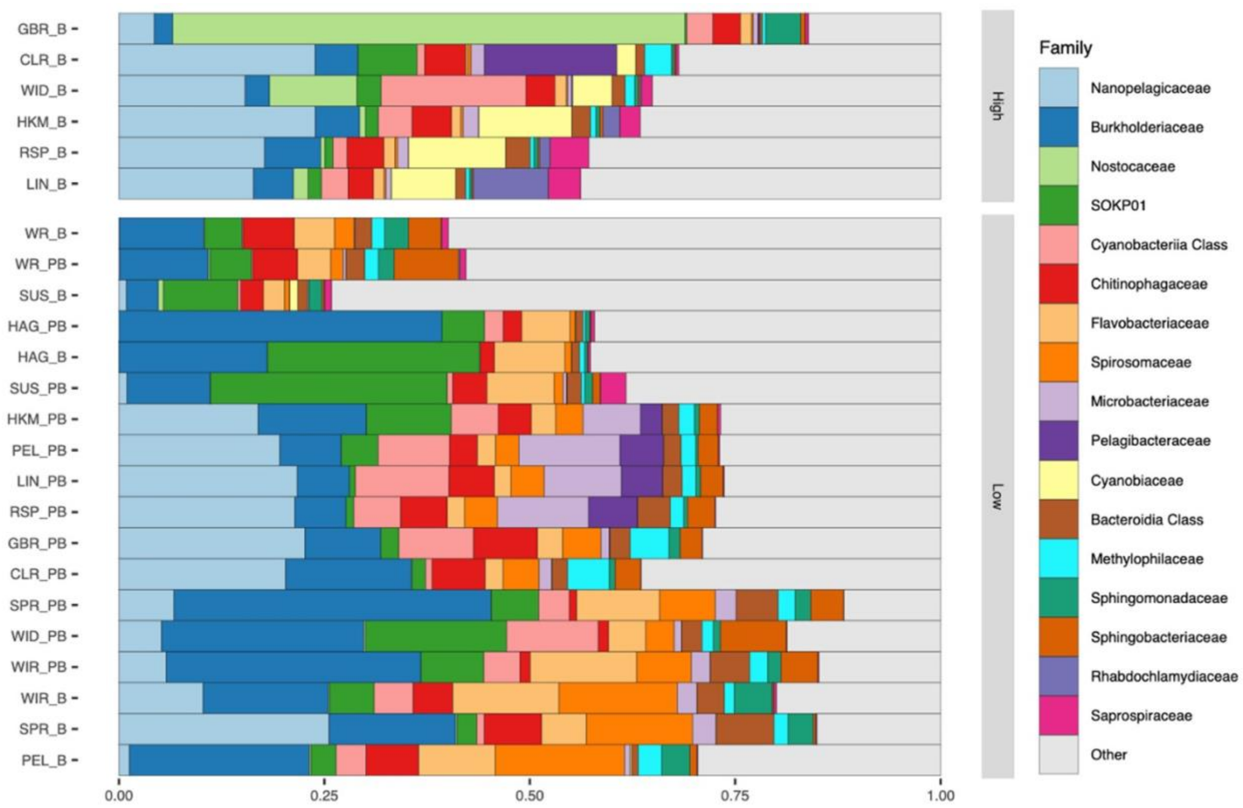


**Figure S5.** Stacked bar plots showing bacterioplankton families of the highest relative abundances based on 16S) and (B mRNA taxonomic annotations. Samples are binned by cyanoHAB abundance. ”PB” = pre-bloom and “B” = bloom.

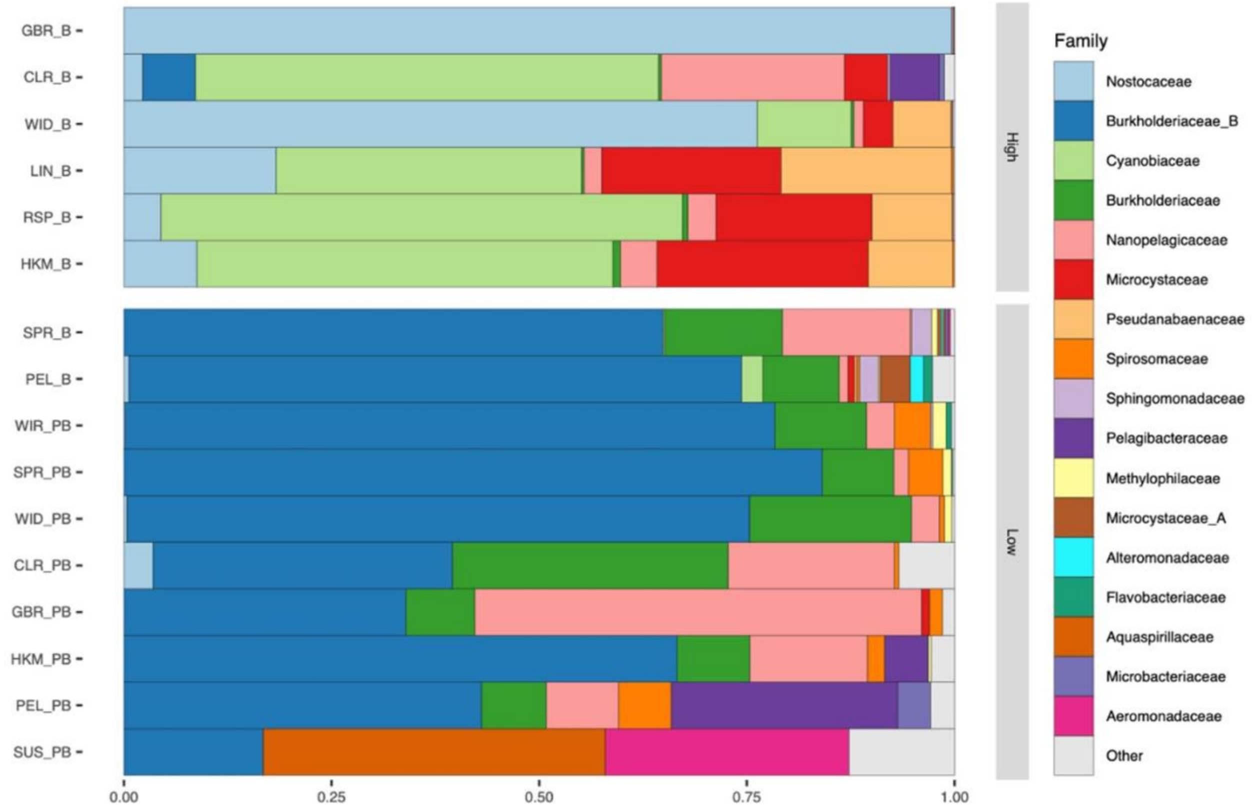


**Figure S6.** Stacked bar plots showing bacterioplankton families of the highest relative abundances based on mRNA taxonomic annotations. Samples are binned by cyanoHAB abundance. ”PB” = pre-bloom and “B” = bloom.

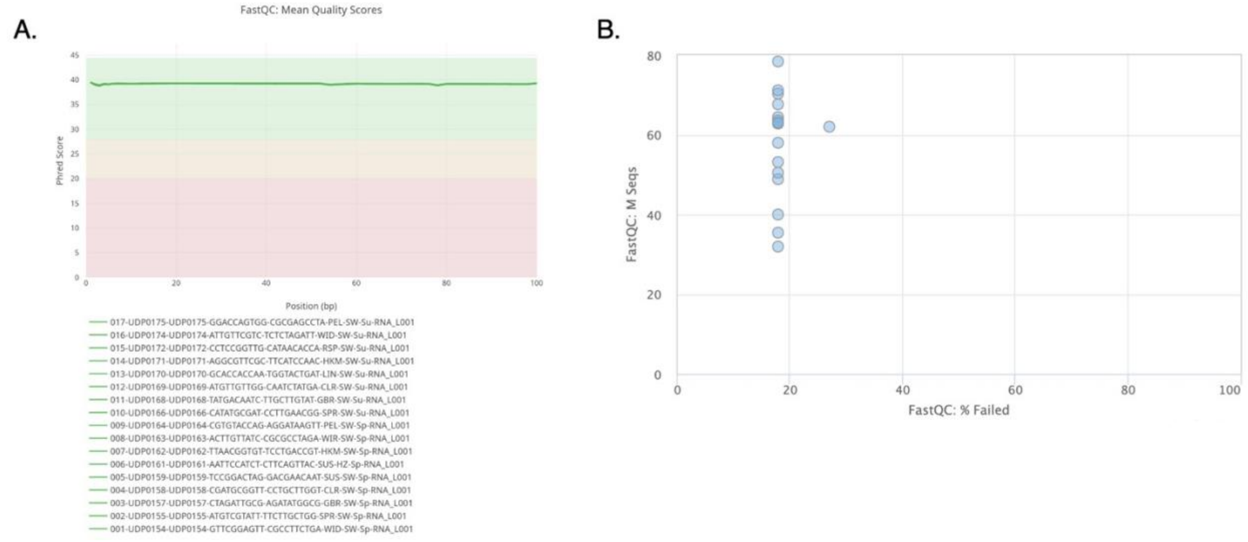


**Figure S7.** mRNA-seq reads displayed high read quality and depth. Read quality reports generated from the CosmosID bioinformatics pipeline displaying the (A) Phred scores of merged forward and reverse samples and (B) read depths and the percent of reads that failed CosmosID quality control steps.

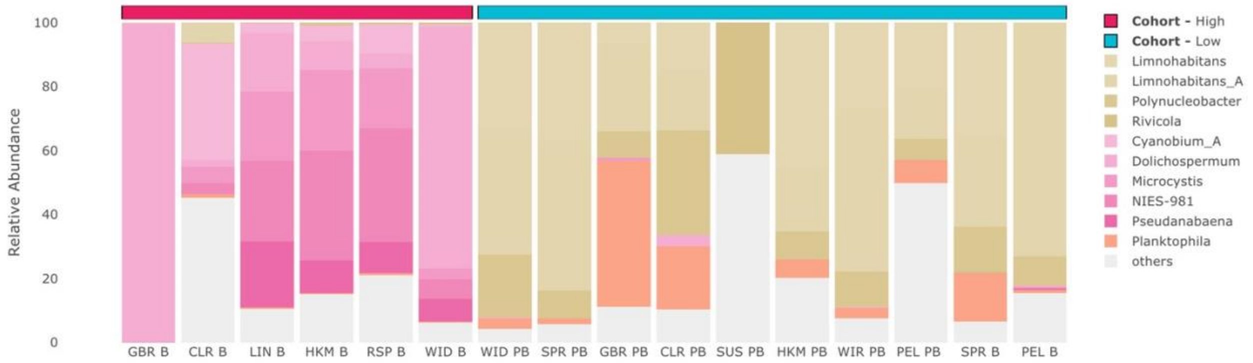


**Figure S8**.  Unique taxa dominate each cyanoHAB abundance bin. Relative abundances of the transcripts from the top 10 most abundant bacterioplankton genera across samples**.** Samples are binned by cyanoHAB abundance (high and low cohorts). This plot was generated by the CosmosID bioinformatics HUB. ”PB” = pre-bloom and “B” = bloom.

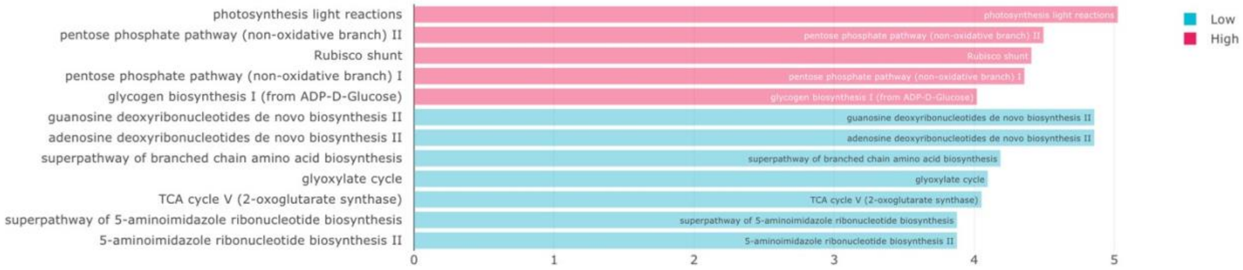


**Figure S9.** MetaCyC pathways differ across sites and between sites of different cyanoHAB impact. Relative abundance (copies per million) of MetaCyc pathways across sample sites binned into high (samples under pink bar) and low (samples under teel bar) cyanoHAB abundnace (or impact). This plot was generated by the CosmosID bioinformatics HUB. ”PB” = pre-bloom and “B” = bloom.

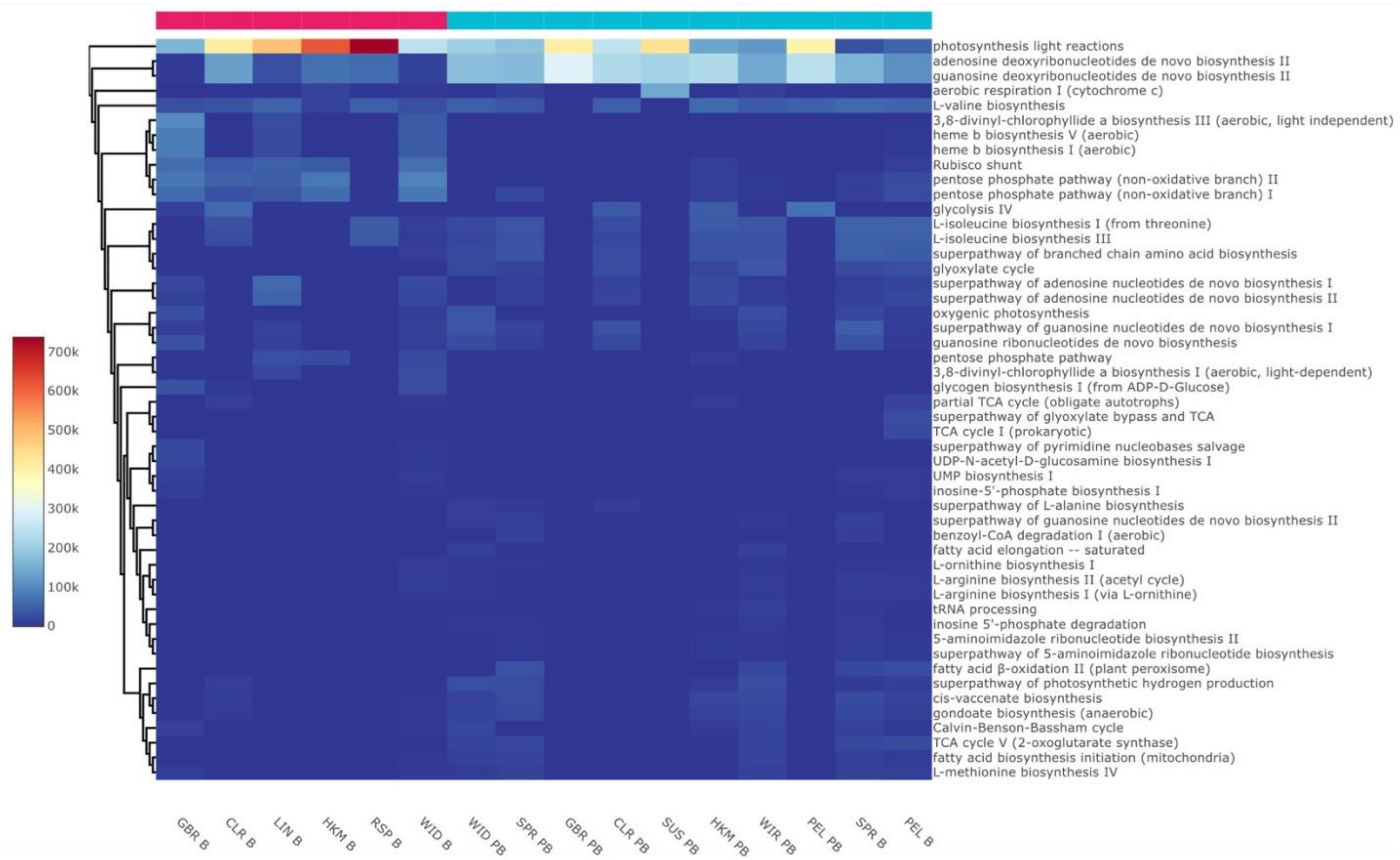


**Figure S10.** MetaCyC pathways differ across sites and between sites of different cyanoHAB impact. Relative abundance (copies per million) of MetaCyc pathways across sample sites binned into high (samples under pink bar) and low (samples under teal bar) cyanoHAB abundance. This plot was generated by the CosmosID bioinformatics HUB. "PB" = pre-bloom and "B" = bloom.

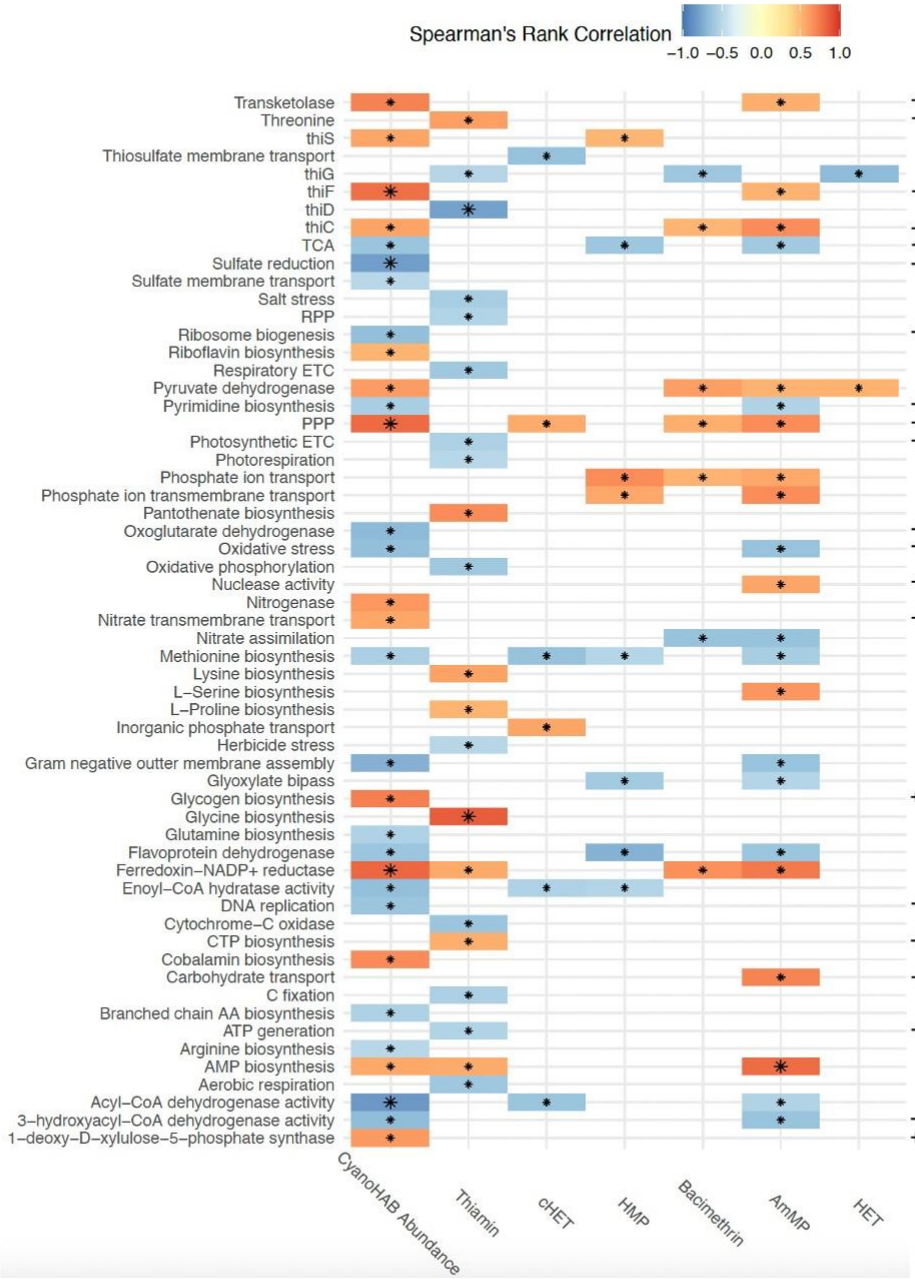


**Figure S11.** Bacterioplankton gene expression that were differentially enriched in cyanoHAB abundance bins also correlated with cyanoHAB abundances (continuous values; cells $ml^{-1}$) and TRC concentrations. Correlogram of GO term and pfam annotations whose CLR-transformed relative abundances (copies per million) significantly correlate (adjusted Spearman $p < 0.05$) with at least one thiamin congener, bacimethrin, and/or cyanoHAB abundance. "+B" = significant lefse enrichment ($p < 0.05$) in high CyanoHAB biomass bin and "-B" = significant lefse enrichment ($p < 0.05$) in low cyanoHAB abundance bin, respectively, based on annotations from at least one CosmosID functional reference database. Threonine = threonine synthase activity; sulfate reduction = assimilatory sulfate reduction; RPP = reductive pentose phosphate pathway; ETC = electron transport chain; PPP = pentose phosphate pathway. Larger asterisks indicate the most significant correlations (adjusted $p < 0.01$).

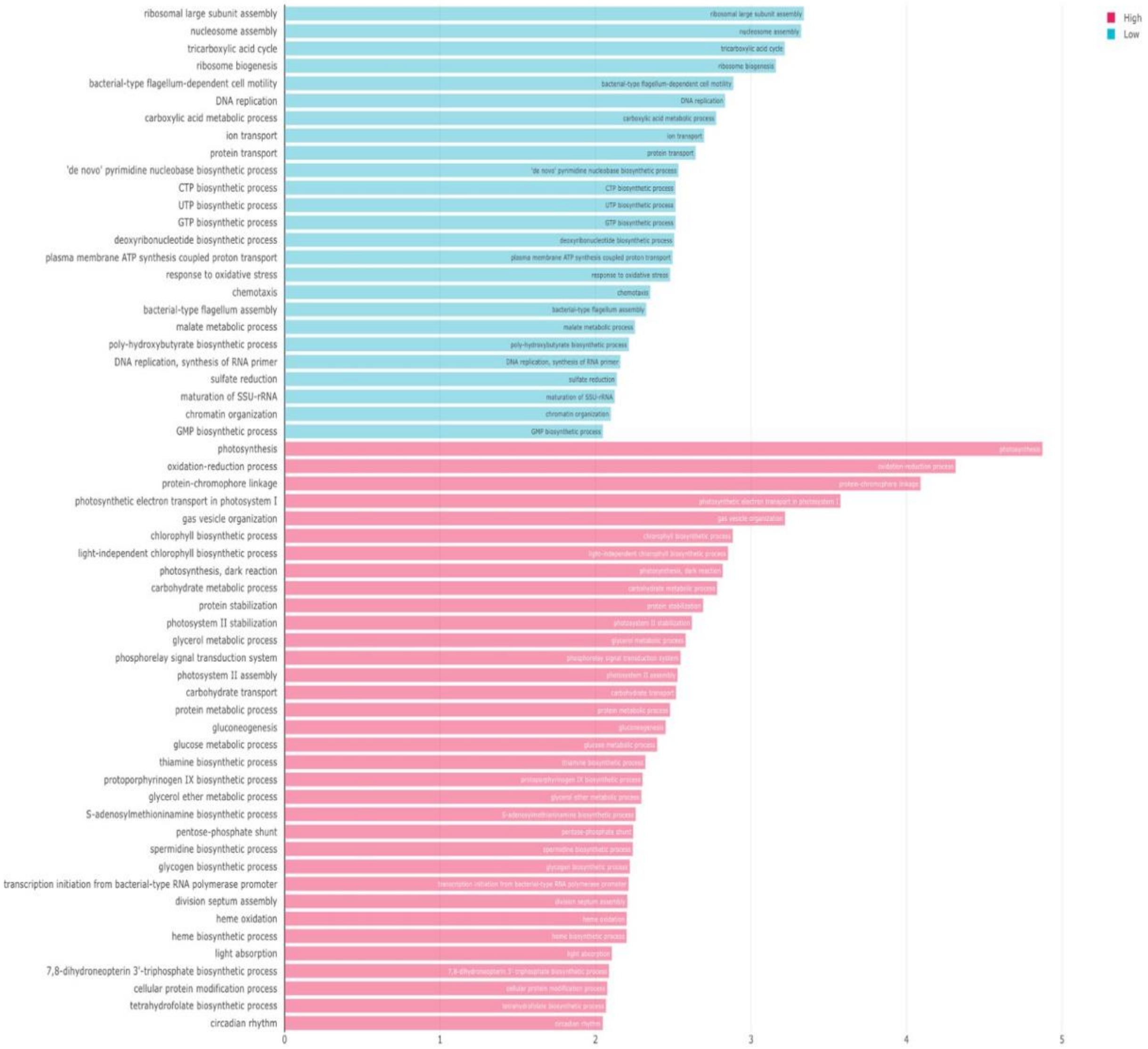


**Figure S12.** GO term-based functional annotations differ based on cyanoHAB abundance. All significant *($p < 0.05$)* GO terms (biological processes) that were enriched in cyanoHAB abundance bins (high and low) based on lefse. The x-axis displays linear discriminant analysis scores. This plot was generated by the CosmosID bioinformatics HUB.

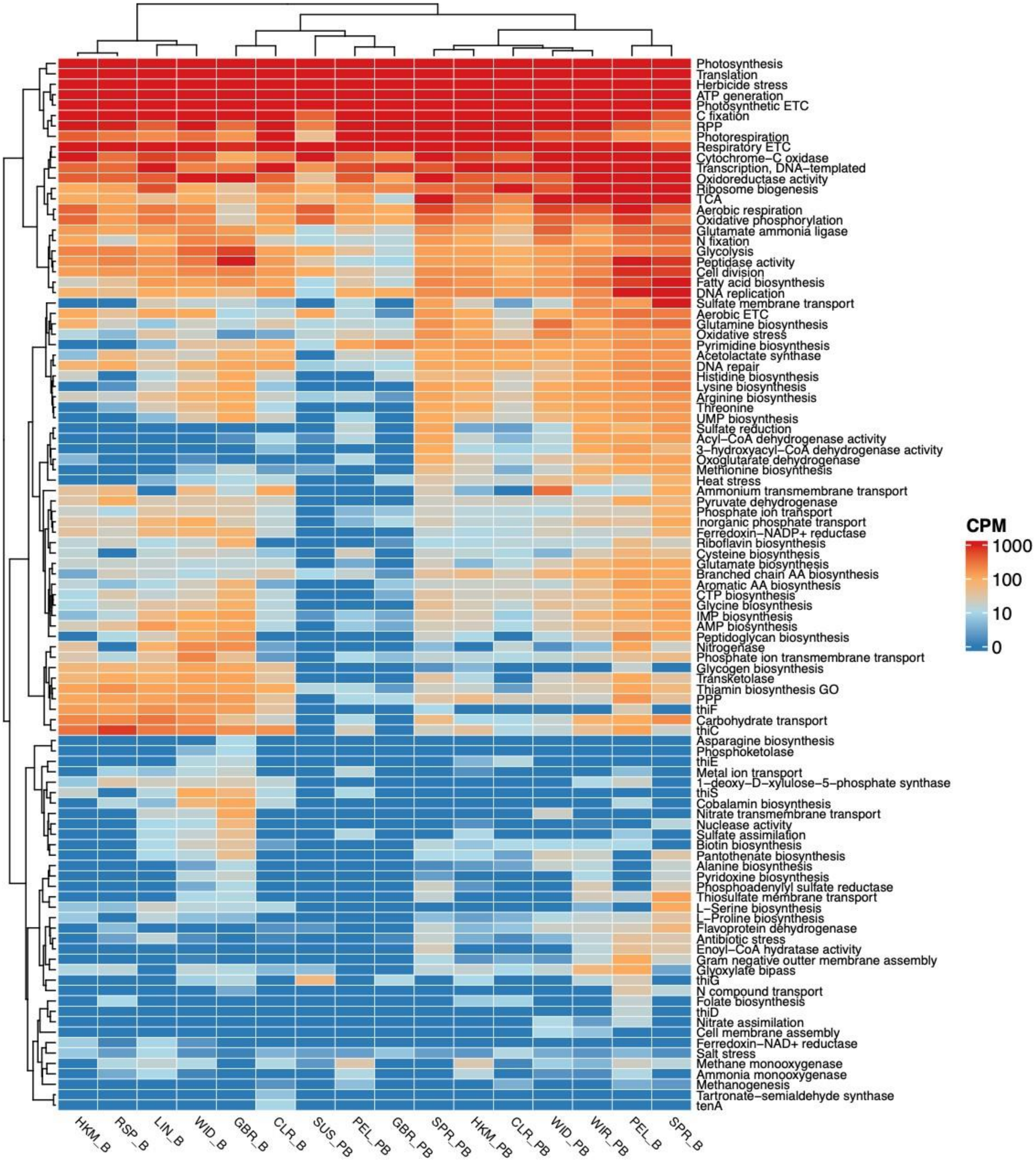


**Figure S13.** All manually curated (see Supp. Methods) functional annotations differ across sample sites. A heatmap of relative abundances (copies per million; CPM) of all the manually curated GO terms and pfam annotations across samples. Sample and function seriation (based on hierarchical clustering) was performed based on Bray-Curtis dissimilarity. ”PB” = pre-bloom and “B” = bloom.
